# Rescue of ribosomal protein bL27 in *Streptococcus pneumoniae* TIGR4 by an alternate protease

**DOI:** 10.64898/2026.08.10.744006

**Authors:** Amarshi Mukherjee, Mohamed O. Nasef, Patrick M. Lindstrom, Nehaal Akavaram, Vipin Chembilikandy, Eriel Martinez, Carlos J. Orihuela, Terje Dokland

## Abstract

*Streptococcus pneumoniae* is a major human respiratory pathogen. The bacterial 70S ribosome is a target of many clinically important antibiotics. The N-terminus of ribosomal protein bL27 extends into the peptidyl transferase center and contributes to the translation process. In Firmicutes, full length bL27 contains an 8-12 amino acid N-terminal extension that is absent from Gram-negative bacteria. This extension is cleaved by the protease Prp, which is absent from organisms lacking the extension. Prp-mediated cleavage of bL27 is essential in *Staphylococcus aureus*, and Prp has been proposed as a potential antibiotic target. Here, we show that in *S. pneumoniae* strain TIGR4, a Δ*prp* mutant remained viable, and produced ribosomes containing cleaved bL27, whereas deletion of *prp* was not tolerated in strain D39. These results suggested the presence of an alternate bL27-processing protease in TIGR4 that was absent from D39. Using a combination of genomics, proteomics and biochemical analyses, we identified this enzyme as the product of previously uncharacterized gene *SP_1145*, encoding a protease that we named Ribosome rescue protease (Rrp). *SP_1145* is carried on a mobile genetic element that is present in strain TIGR4, but absent from D39. Our findings shed light on an alternative mechanism for bL27 maturation, and indicate that some strains of *S. pneumoniae* harbor horizontally acquired redundant pathways for this essential ribosome processing step.

## INTRODUCTION

The bacterial ribosome is a prime target for numerous antibiotics [1–4], many of which are in use against clinically important bacteria such as *Staphylococcus aureus* and *Streptococcus pneumoniae* [5–8]. The bacterial ribosome (70S) consists of a large (50S) and a small (30S) subunit, and is composed of 54 proteins and three RNAs [9, 10]. The catalytic steps in the translation process are carried out primarily by the RNA; however, the protein components are necessary for correct folding and assembly of the ribosome and may play roles in modulating its activity [9, 11]. bL27, product of the *rpmA* gene, is a ribosomal large subunit protein that does not have an equivalent in eukaryotes and archaea [12]. bL27 is uniquely positioned in the ribosome where its N-terminus reaches into the peptidyl transferase center and interacts with the tRNA [13]. However, its involvement in peptide bond formation or other aspects of the translation process is still unclear [14, 15]. An *Escherichia coli* Δ*rpmA* mutant was viable, but exhibited greatly impaired growth, reduced peptidyl transferase activity and accumulation of 40S ribosomal large subunit precursors [12, 16].

In the Firmicutes, which includes streptococci and staphylococci as well as listeriae, bacilli, and clostridiae, bL27 has an additional, conserved 8-12-residue N-terminal extension that is not present in bL27 from Gram-negative organisms such as *E. coli* [17, 18](Fig. 1A). This N-terminal extension is not observed in structures of Gram-positive ribosomes [19–21]. We previously described a cysteine protease that cleaves the N-terminal extension of bL27 in *Staphylococcus aureus* [18]. This protease is encoded by a gene that is located between *rplU* (encoding bL21) and *rpmA* in the same operon (Fig. 1B). Because we originally identified this protein for its role in cleavage of bacteriophage proteins, we called it Prp, for “<u>P</u>hage-related ribosomal <u>p</u>rotease” [18]. The *prp* gene is present in the genomes of all Firmicutes and other bacteria that encode bL27 with the N-terminal extension, but not in *E. coli* and other bacteria that lack the N-terminal extension [18]. Both the presence of the N-terminal extension of bL27 and its ability to be cleaved were found to be essential for normal ribosome assembly in *S. aureus* using an expression system in which the wildtype bL27 could be turned off and replaced with either a truncated or an uncleavable version of bL27 [22]. The role of the N-terminal extension of bL27 in Gram-positive bacteria is so far unknown.

**Figure 1.**
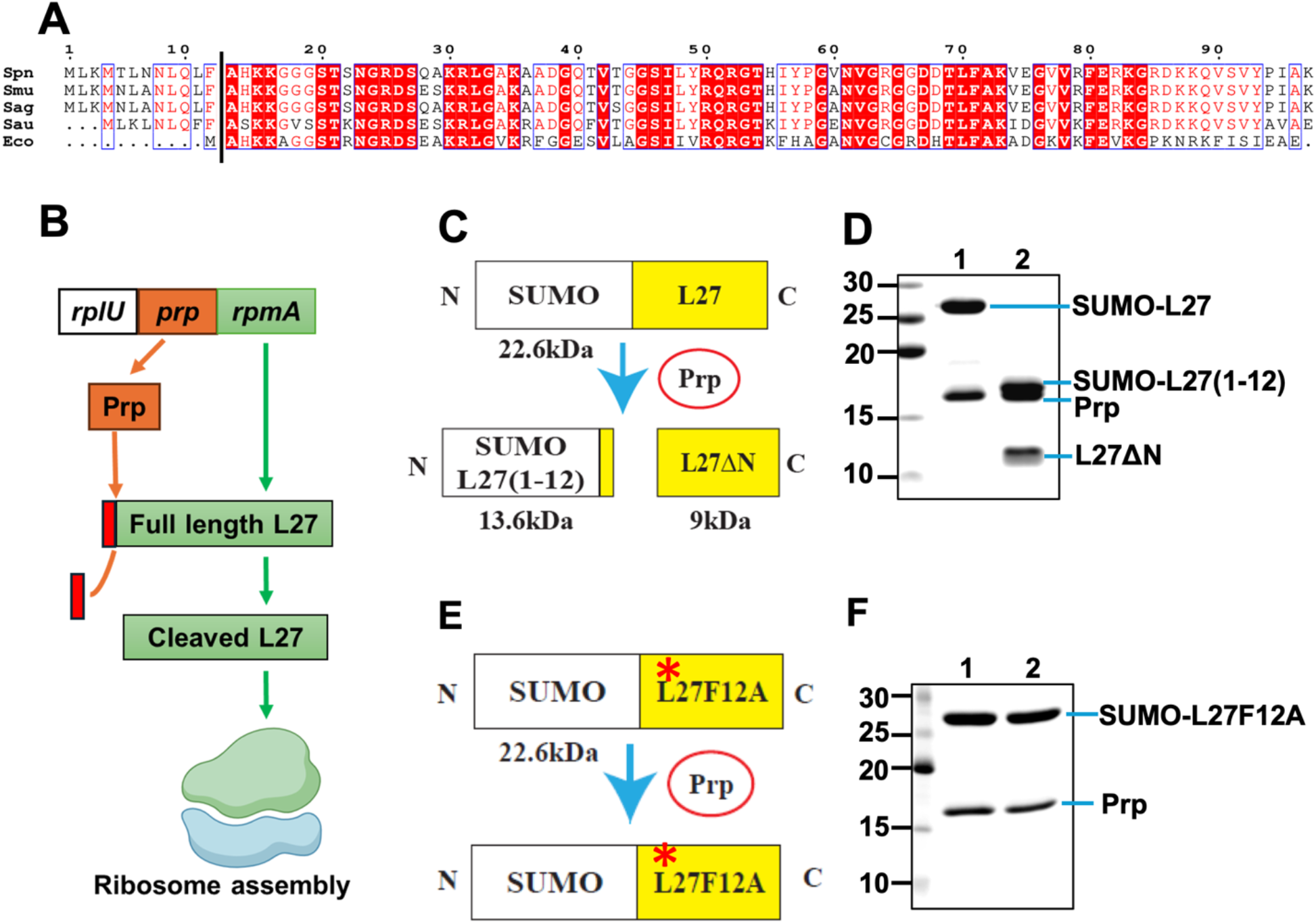
In vitro cleavage of bL27. **(A)** Aligned amino acid sequences of bL27 proteins from *S. pneumoniae* (Spn), *S. mutans* (Smu), *S. agalactiae* (Sag), *S. aureus* (Sau), and *E. coli* (Eco). The F|A cleavage site is shown as a vertical line. **(B)** Schematic representation of bL27 cleavage by Prp and its incorporation into ribosomes. **(C)** Schematic representation of the in vitro cleavage of SUMO-L27 by Prp. **(D)** Analysis of SUMO-L27 digestion by Prp. Equimolar amounts of SUMO-L27 and Prp were mixed, immediately boiled in Laemmli sample buffer, and loaded onto SDS-PAGE (lane 1). An identical sample was incubated in assay buffer at 37 °C for 1 h before boiling (lane 2). Molecular weights of markers are indicated (kDa). **(E)** Schematic representation of the lack of cleavage of SUMO-L27F12A by Prp. **(F)** Analysis of SUMO-L27F12A digestion by Prp as described for (D).

*S. pneumoniae* (pneumococcus) is one of the most important respiratory pathogens in humans [5–7], responsible for at least half a million deaths among children worldwide every year [23], and a common cause of bacterial meningitis [24]. Development of new antibiotics has not kept up with the increase in antibiotic resistance in *S. pneumoniae* and other bacteria [25–27]. Prp has previously been proposed as a target for novel antibiotics against *S. aureus* [18, 22, 28, 29]. However, in this study we show that *prp* is not absolutely essential in *S. pneumoniae* strain TIGR4. A *S. pneumoniae* TIGR4 Δ*prp* mutant remained viable; ribosomes isolated from a TIGR4 Δ*prp* mutant contained processed Prp, and a TIGR4 Δ*prp* cell lysate was able to cleave recombinant bL27 in vitro. In contrast, *prp* appeared to be essential in *S. pneumoniae* strain D39. Taken together, these results indicated the presence of an alternate protease that could carry out the processing of bL27 in TIGR4. Using a combination of genomics, proteomics and biochemistry, we have shown that the alternate protease is the product of TIGR4 gene *SP_1145*, encoding a previously uncharacterized protease belonging to the Ntn hydrolase family that includes the eukaryotic proteasome α and β subunits and the bacterial Anbu protein [30, 31]. Moreover, the *SP_1145* gene appears to be part of a horizontally acquired mobile genetic element that is absent from D39. We named this alternate protease Rrp for “<u>R</u>ibosome rescue <u>p</u>rotease.” Our results imply that cleavage of the N-terminal extension of bL27 is indeed essential in *S. pneumoniae* and other Gram-positive bacteria, but that some strains of *S. pneumoniae* harbor redundant pathways capable of carrying out the essential bL27 processing and thereby rescue ribosome assembly in the absence of Prp.

## RESULTS

### 1. Cleavage of bL27 by Prp in vitro

To examine the *in vitro* activity of Prp and its cleavage of bL27, we cloned *prp* from *S. pneumoniae* TIGR4, a virulent serotype 4 strain (Table 1) into the pET21a vector with the addition of a C-terminal myc-His_6_ tag, and expressed the protein in *E. coli*. We also expressed full-length TIGR4 bL27 as an N-terminally His_6_-tagged SUMO fusion protein , referred to here as SUMO-L27. Both proteins were purified to homogeneity by Ni-NTA affinity and size exclusion chromatography (SEC). bL27 is highly conserved with 97% amino acid identity across streptococcal species (Fig. 1A). Prp is more variable, with only around 50% identity between streptococcal species, although the active site catalytic dyad is completely conserved [18].

**Table 1.**
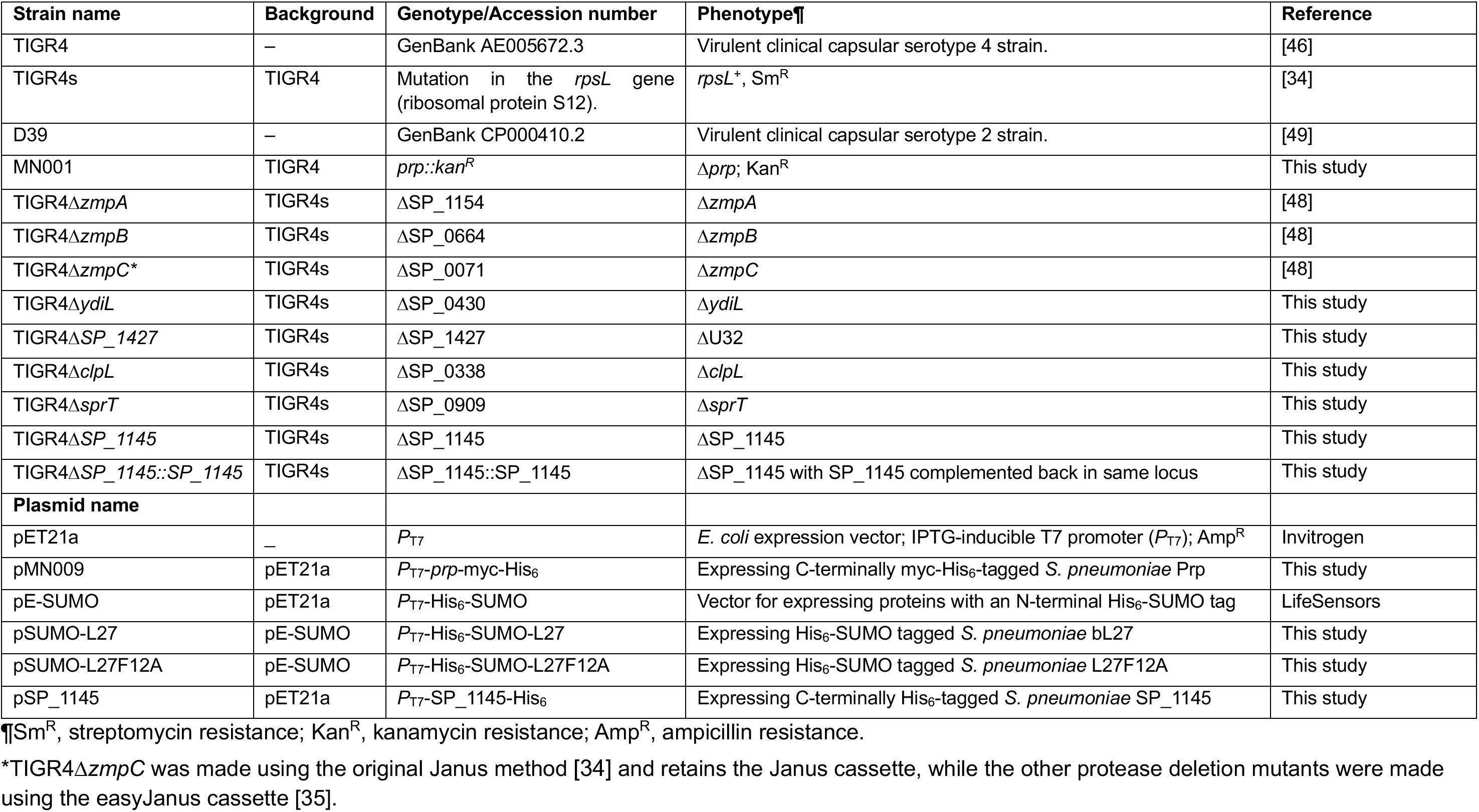
Strains and plasmids used in this study.

Cleavage of the 22.6 kDa SUMO-L27 fusion protein at the expected Prp cleavage site between Phe 12 and Ala 13 predicted by sequence comparison to *S. aureus* (Fig. 1A) would yield two bands of 13.6 kDa and 9.0 kDa, corresponding to the SUMO moiety fused to the first 12 amino acids of bL27 [SUMO– L27(1–12)] and the mature bL27 protein (L27ΔN; residues 13–97), respectively (Fig. 1C). When SUMO-L27 was mixed with an equimolar amount of TIGR4 Prp, it was fully cleaved after 1 h at 37 °C (Fig. 1D). Full-length mass spectrometry identified a fragment of 9,014 Da, corresponding to the expected size of L27ΔN (9,014.15 Da; Fig. S1).

We also generated a SUMO-L27 construct with a Phe 12 to Ala mutation (F12A) in the predicted F|A cleavage site (SUMO-L27F12A, Fig. 1E), which had previously been shown to block cleavage by Prp in *S. aureus* [18]. SUMO-L27F12A was expressed and purified in the same way as the wildtype SUMO-L27. When this protein was mixed with Prp, no cleavage occurred (Fig. 1F), consistent with the expected sequence specificity of Prp and our previously reported results in *S. aureus* [18].

### 2. The N-terminal extension of bL27 is cleaved in *S. pneumoniae* in the absence of Prp

To investigate the role of Prp *in vivo*, we used homologous recombination to produce a *prp* knockout strain (Δ*prp*) in *S. pneumoniae* TIGR4, where *prp* (TIGR4 gene *SP_1106*) was replaced with a kanamycin resistance cassette (gene *aphIII*), producing the kanamycin resistant (Kan^R^) strain MN001 (Table 1, Fig. 2A). Surprisingly, this mutant was viable, albeit with delayed growth, displaying a twofold increase of the lag phase time compared to the TIGR4 wildtype (Fig. 2B). During exponential growth, the Δ*prp* mutant had a similar generation time (63 min) to the wildtype (61 min); however, the Δ*prp* mutant never reached the same cell density as TIGR4 (Fig. 2B). Whole-genome sequencing confirmed that the *prp* gene had been replaced by *aphIII* in its original locus.

**Figure 2.**
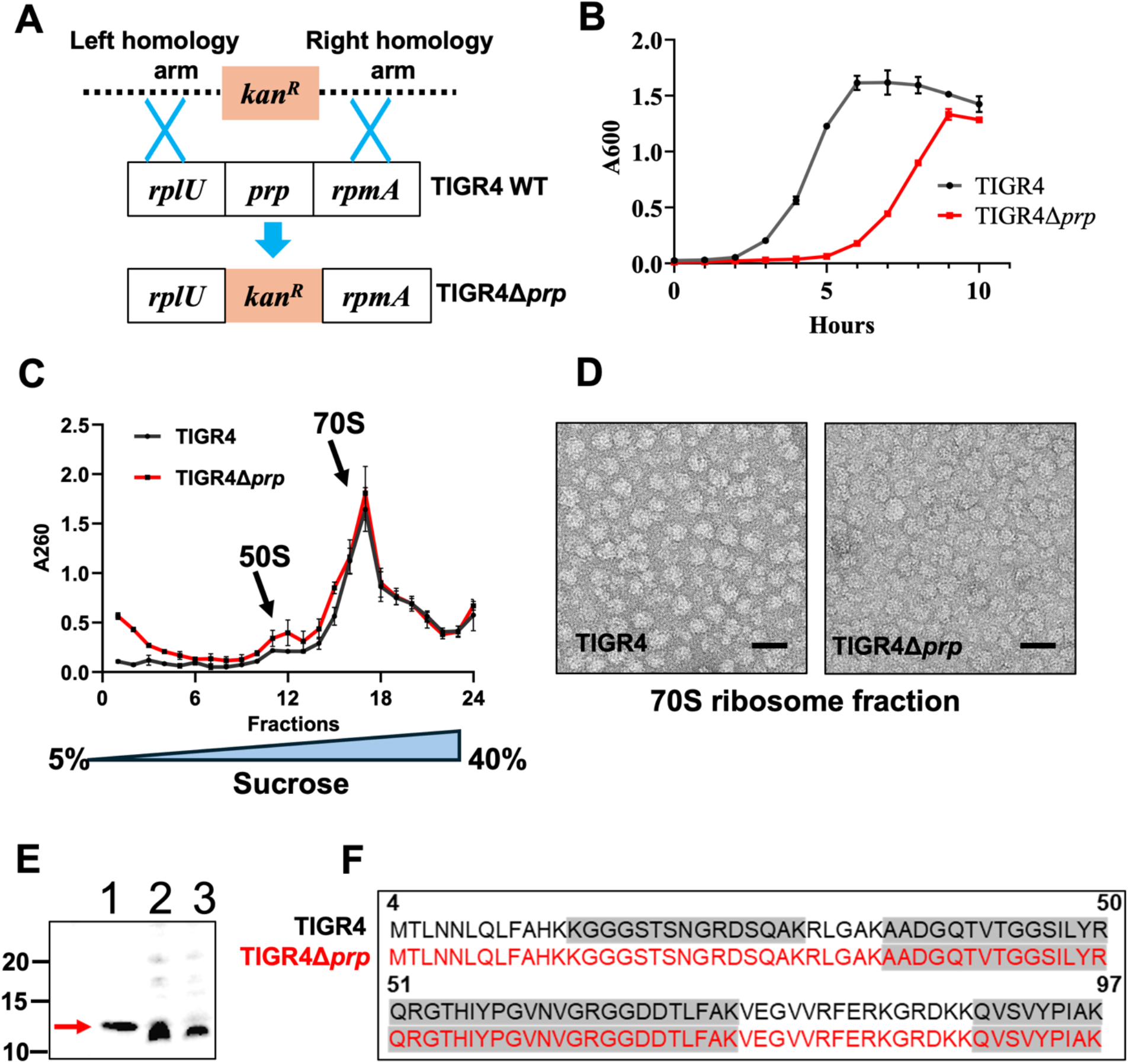
Characterization of *prp* mutant strains. **(A)** Schematic representation of *prp* deletion in TIGR4. **(B)** Growth curve for the TIGR4 Δ*prp* mutant (strain MN001; red curve) compared to the TIGR4 wildtype strain (black curve). **(C)** Fractionated sucrose gradient profile for ribosomes purified from TIGR4 wildtype (black curve) and TIGR4 Δ*prp* (red curve) strains. **(D)** Negative stain EM of purified 70S ribosomes from TIGR4 wildtype (WT) and TIGR4 Δ*prp* strains. Scale bar = 50 nm. **(E)** Western blot analysis of ribosomes using an anti-*S. aureus* bL27 antibody. Lane 1, SUMO-L27 treated with purified Prp; Lane 2, purified ribosomes from *S. pneumoniae* TIGR4 wildtype; Lane 3, purified ribosomes from the TIGR4 Δ*prp* strain. The band corresponding to the cleaved L27ΔN is indicated by the red arrow. **(F)** Mass spectrometry analysis of bL27 in TIGR4 wildtype (black) and TIGR4 Δ*prp* (red) ribosomes. Peptides detected above 80% confidence level are highlighted in gray.

In light of observations in *S. aureus*, where overexpression of inactive Prp mutants (PrpC34A and PrpC34S) had a dominant negative phenotype and led to accumulation of 40S ribosomal complexes [18], we hypothesized that the *S. pneumoniae* Δ*prp* mutant would be defective in ribosome assembly. We isolated ribosomes from the TIGR4 wildtype strain and the Δ*prp* mutant strain at mid-log phase (A_600_=0.6 OD), according to established procedures [19, 32]. The ribosomes were separated on 5–40% sucrose gradients in 50 mM NH_4_Cl in the presence of 15 mM Mg^2+^, where 70S ribosomes are stable. The sucrose gradient profile was similar in both cases, showing the expected peaks corresponding to 50S subunits and 70S ribosomes, with a slight increase in the 50S peak in the Δ*prp* strain compared to the wildtype (Fig. 2C). The purified 70S ribosomes from WT and Δ*prp* strains appeared normal by electron microscopy (Fig. 2D).

To check for the presence or absence of uncleaved bL27 in the ribosomes, we separated purified wildtype and Δ*prp* ribosomes by SDS-PAGE and did a Western blot with a polyclonal anti-bL27 antibody that we originally raised against *S. aureus* bL27 [18], but that also works for *S. pneumoniae* (Fig. 2E). SUMO-L27 cleaved with Prp was used as a control. Surprisingly, there was no difference in the mobility of bL27 between the wildtype and Δ*prp* ribosomes (Fig. 2E), suggesting that the bL27 protein had been cleaved even in the absence of Prp. When the band corresponding to bL27 was excised and analyzed by mass spectrometry (MS), no peptides corresponding to the N-terminal extension of bL27 were detected at a confidence level >80% (Fig. 2F, Table S1), further indicating that bL27 cleavage occurs in the Δ*prp* mutant. Prp was not detected by MS in either the TIGR4 wildtype or Δ*prp* mutant (Table S2) These results suggested the presence of another protease capable of cleaving bL27 in *S. pneumoniae* TIGR4.

### 3. bL27 is cleaved in *S. pneumoniae* cell extracts

To further investigate the cleavage of bL27 by *S. pneumoniae*, we developed an in vitro cleavage assay in which the purified SUMO-L27 was mixed with bacterial cell lysates, incubated for 3 h at 37 °C, separated by SDS-PAGE, and probed by Western blotting using either an anti-His_6_ antibody or the anti-bL27 antibody (Fig. 3A). As a positive control, we used a lysate from the Prp-expressing *E. coli* strain. The amount of SUMO-L27 digestion was quantified relative to the 0 h reaction by densitometric scan of the bands corresponding to the remaining uncleaved SUMO-L27.

**Figure 3.**
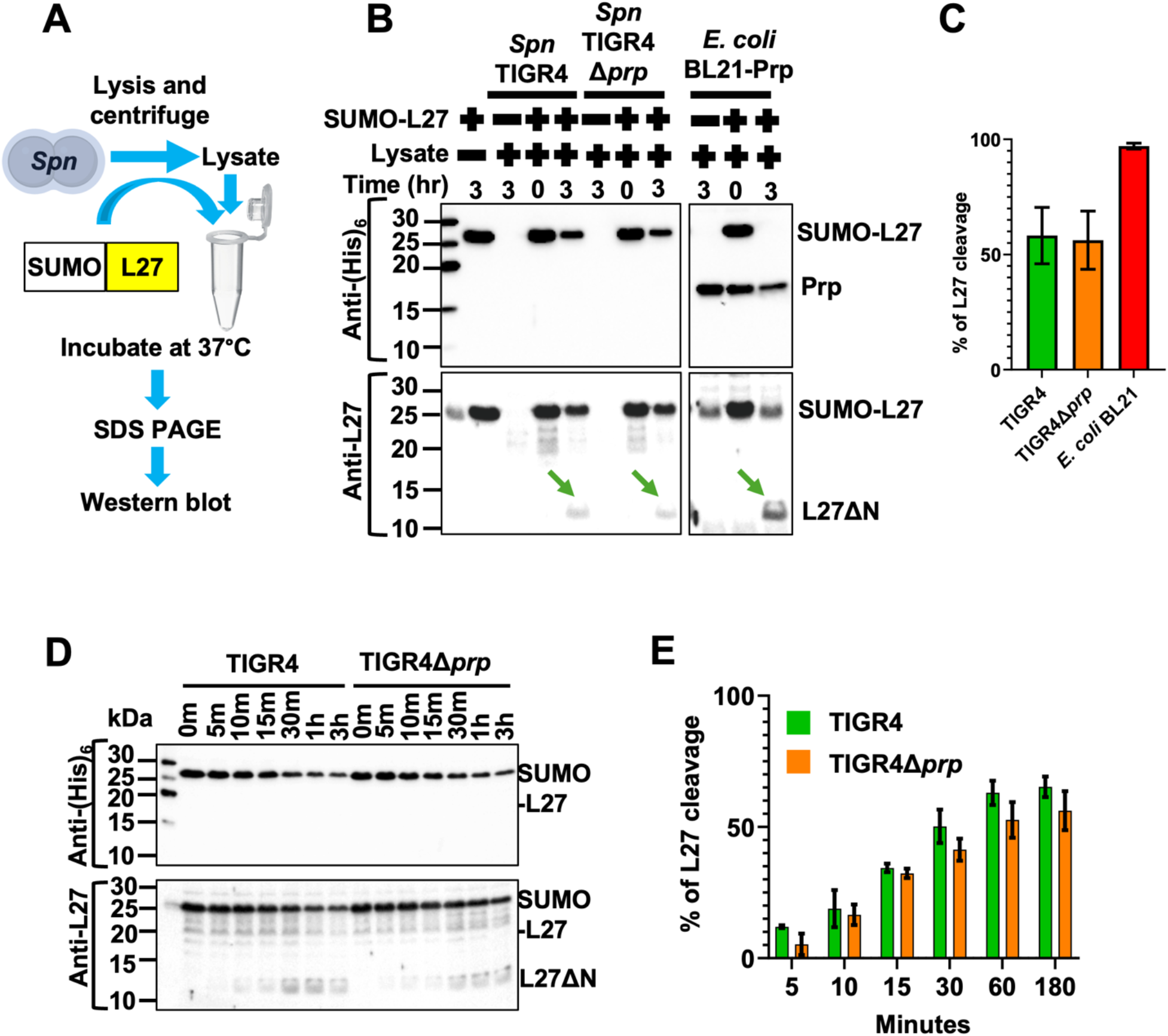
Cleavage of bL27 by cell extracts. **(A)** Schematic representation of the SUMO-L27 cleavage assay by *S. pneumoniae* cell lysates. **(B)** In vitro cleavage of SUMO-L27 by lysates of *S. pneumoniae* TIGR4 and TIGR4 Δ*prp* for 3 h at 37 °C. The 0 h sample was prepared by adding Laemmli sample buffer immediately after mixing, followed by boiling for 5 min. As a positive control for SUMO-L27 cleavage, an *E. coli* BL21 lysate expressing His_6_-tagged Prp (BL21-Prp) was used. Samples were separated by 15% SDS-PAGE and analyzed by immunoblotting with anti-His_6_ and anti-*S. aureus* bL27 antibodies. Bands corresponding to uncleaved SUMO-L27, Prp and cleaved L27ΔN (green arrows) are labeled. **(C)** Quantitation of SUMO-L27 cleavage, measured as percentage reduction of the SUMO-L27 signal (average of both Western blots) from (B) at 3 h compared to 0 h. **(D)** Kinetic assay of SUMO-L27 cleavage by TIGR4 and TIGR4 Δ*prp* cell lysates, incubated for up to 3 h, separated by SDS-PAGE and analyzed by immunoblotting with anti-His_6_ or anti-bL27 antibodies, as above. **(E)** Quantitation of SUMO-L27 cleavage by TIGR4 wildtype (green) and TIGR4 Δ*prp* (orange) as a function of time, measured as reduction in SUMO-L27 signal from (D), averaged from both blots.

This assay showed that the SUMO-L27 fusion protein was about 57±10% cleaved after 3 h in the presence of either the wildtype TIGR4 or the TIGR4 Δ*prp* lysate, observed as a weakening of the SUMO-L27 bands in the Western blots (Fig. 3B, C). Although the cleaved SUMO-L27(1-12) fragment was not detected with the anti-His_6_ antibody, a band corresponding to the cleaved bL27 (L27ΔN) could be detected when probed with the anti-bL27 antibody (Fig. 3B). Importantly, there was no difference in cleavage between the TIGR4 wildtype and Δ*prp* lysates (Fig. 3B, C), demonstrating that a protease present in the TIGR4 strain was able to cleave bL27 in the absence of Prp. By comparison, the Prp-expressing *E. coli* lysate (BL21-Prp) caused >95% cleavage of SUMO-L27 after 3 h of incubation (Fig. 3B, C), whereas no cleavage of SUMO-L27 occurred in an *E. coli* lysate in the absence of Prp expression (Fig. S2). A time-course experiment in which SUMO-L27 was mixed with wildtype and Δ*prp* lysates for up to 3 h showed that SUMO-L27 was cleaved efficiently by the TIGR4 Δ*prp* lysate, but with a delayed onset compared to the wildtype (Fig. 3D, E).

The SUMO-L27F12A mutant protein was then subjected to the same cleavage assay. We already showed that the SUMO-L27F12A mutant could not be cleaved by purified Prp (Fig. 1F). Surprisingly, the SUMO-L27F12A mutant protein was cleaved, albeit less efficiently than SUMO-L27, by both the TIGR4 wildtype and TIGR4 Δ*prp* lysates (Fig. 4A, B), suggesting that the presumptive alternate protease had a distinct cleavage specificity from Prp.

**Figure 4.**
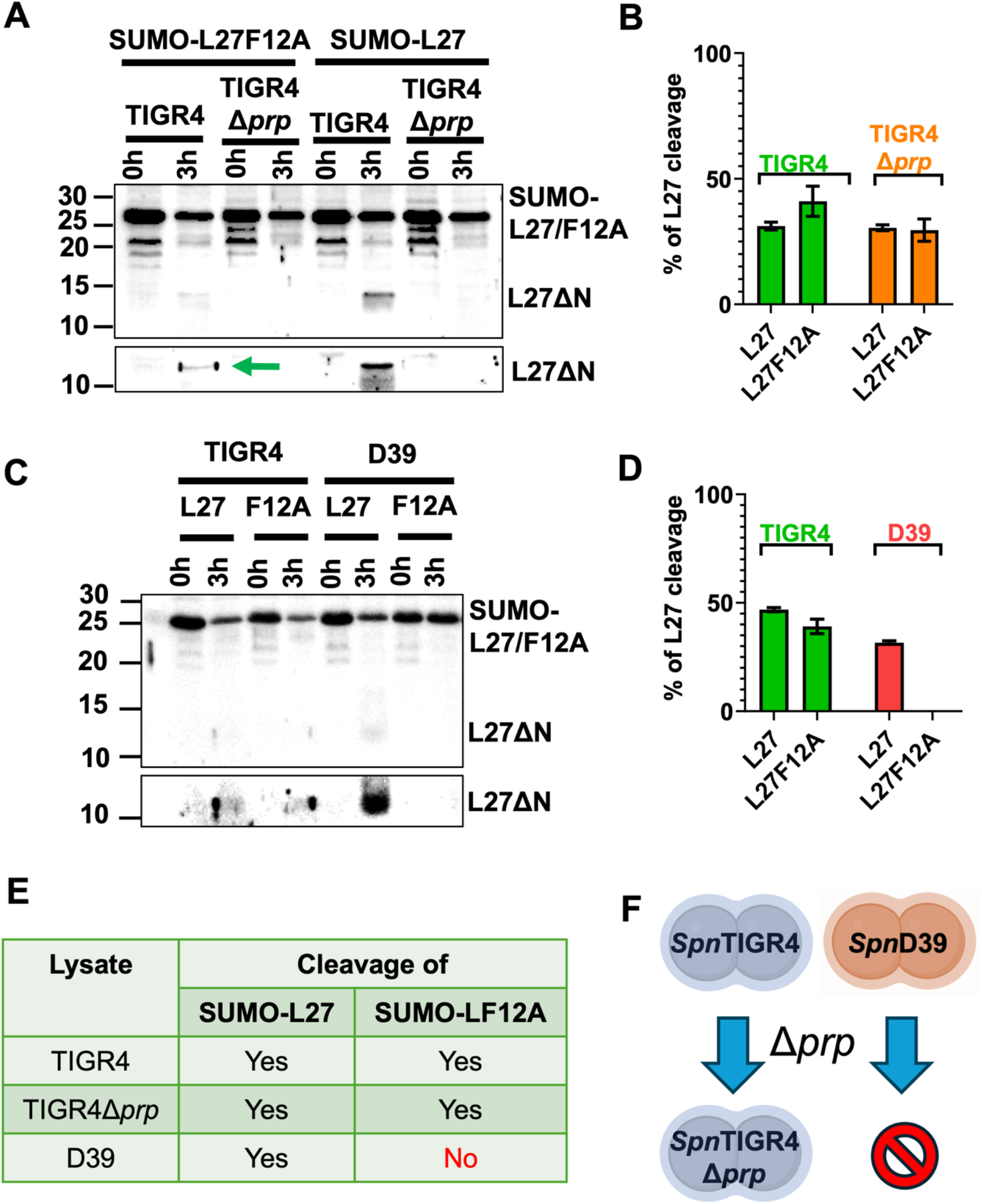
Cleavage of the L27F12A mutant by cell extracts. **(A)** Cleavage of SUMO-L27 and SUMO-L27F12A by TIGR4 wildtype and TIGR4 Δ*prp* cell lysates, separated by SDS-PAGE and detected by anti-bL27 antibody. A higher exposure of the Western blot, showing the L27ΔN band from SUMO-L27F12A more clearly (green arrow) is shown below. **(B)** Densitometric quantitation of cleavage of SUMO-L27 and SUMO-L27F12A by the TIGR4 wildtype (green) and TIGR4 Δ*prp* (orange) strains from the Western blot in (A). **(C)** Cleavage of SUMO-L27 and SUMO-L27F12A by lysates of *S. pneumoniae* strains TIGR4 and D39, separated by SDS-PAGE and detected with the anti-bL27 antibody. A higher exposure of the blot is shown below. **(D)** Densitometric quantitation of the Western blot in (C). **(E)** Table summary of SUMO-L27 and SUMO-L27F12A cleavage by cell lysates. **(F)** Schematic diagram illustrating the result of deleting *prp* in TIGR4 and D39 strains.

We then tested whether SUMO-L27F12A could be cleaved by a lysates of *S. pneumoniae* D39, a virulent serotype 2 strain. Strikingly, while the SUMO-L27 wild-type protein was cleaved efficiently by both D39 and TIGR4, the SUMO-L27F12A mutant was only cleaved by TIGR4 and not by D39 (Fig. 4C, D). This result indicated that the presumptive alternate protease capable of cleaving both SUMO-L27 and SUMO-L27F12A was present in TIGR4, but absent from D39 (Fig. 4E). Consistent with this result, attempts at deleting the *prp* gene in strain D39 were not successful (Fig. 4F).

### 4. Screening for the alternate protease in *S. pneumoniae* TIGR4

To search for the alternate protease, we compared the annotated and publicly available genomes of TIGR4 (GenBank ID AE005672) and D39 (CP000410). According to published literature, *S. pneumoniae* encodes 34 known proteases [33]. Of these 34, the genes encoding CbpG (*SP_0390*), PrsW (*SP_0026*), pyroglutamyl peptidase II (Pcp2; *SP_2060*), and YdiL (*SP_0430*) are frameshifted in D39, while gene *SP_0071*, encoding Zinc metalloprotease ZmpC, is absent from D39, thus constituting possible candidates for the alternate protease. However, given that the protease of interest was presumed to be cytosolic, we excluded CbpG, a surface-exposed protein, PrsW, which is involved in cell membrane stress response, and Pcp2, which has a highly specific enzymatic function [33].

Using Janus replacement [34, 35] (Fig. S3), we generated deletion mutants of *SP_0430* (Δy*diL*), *SP_1429* (ΔU32), and *SP_0071* (Δ*zmpC*) in the TIGR4s background (Table 1). (TIGR4s is a derivative of TIGR4 with a mutation in the *rpsL* gene, conferring streptomycin resistance, used for the Janus cassette replacement procedure [34].) In addition, we tested four other previously described mutants of protease-encoding genes: *SP_1154* (Δ*zmpA*; encoding ZmpA, an IgA1 protease), *SP_0664* (Δ*zmpB*; encoding ZmpB, a zinc metalloprotease), *SP_0338* (Δ*clpL*; encoding ClpL, a manganese-dependent chaperone-like protease), and *SP_0909* (Δ*sprT*; encoding SprT, a zinc metalloprotease).

Lysates from each mutant strain were prepared and tested for their ability to cleave SUMO-L27 and SUMO-L27F12A using the assay described above. If a given protease were responsible for cleaving the SUMO-L27F12A mutant, then deletion of that protease should abolish cleavage of SUMO-L27F12A in the corresponding lysate. As a control, a D39 lysate, which is unable to cleave SUMO-L27F12A, was included. However, no differences in cleavage activity were observed between any of the mutant lysates and the TIGR4s wildtype strain for either substrate (Fig. 5A). These results suggested that the alternate protease responsible for this activity was likely a protein that had not yet been characterized.

**Figure 5.**
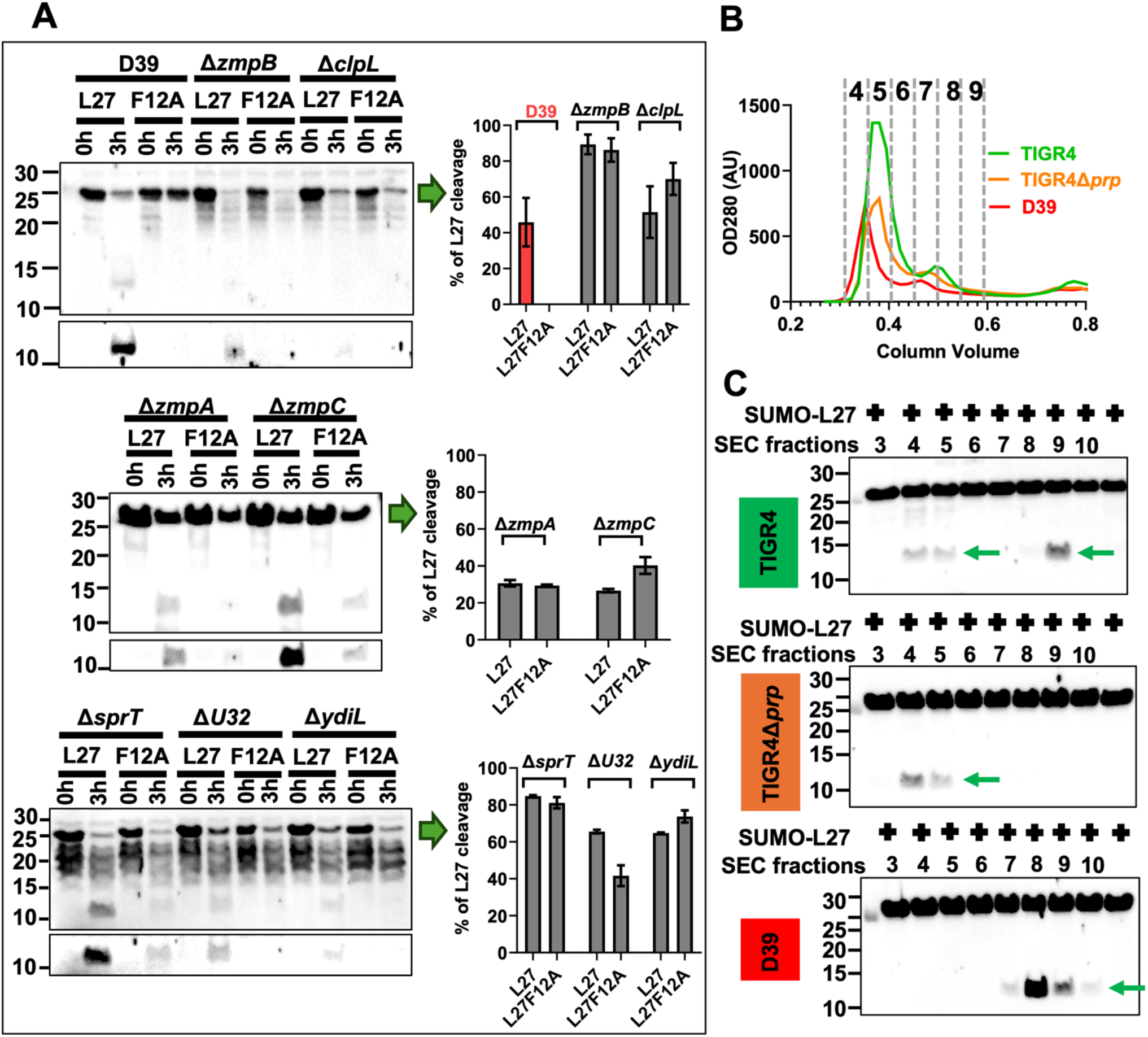
Cleavage assay and protein fractionation by SEC. **(A)** Cleavage assay performed with different protease deletion mutants, probed with the anti-bL27 antibody, followed by densitometric quantitation. A D39 lysate (red bar) was used as a control for cleavage of SUMO-L27 and lack of cleavage of SUMO-L27F12A. **(B)** SEC separation of cell lysates of TIGR4 (green), TIGR4 Δ*prp* (orange) and D39 (red) strains. Fractions collected are indicated by the numbers. **(C)** Cleavage of SUMO-L27 by SEC fractions from (B), separated by SDS-PAGE and probed with the anti-bL27 antibody. Bands corresponding to the cleaved L27ΔN are indicated by green arrows.

### 5. Biochemical identification of the unknown protease

To home in on the unknown protease, we used a biochemical approach in which lysates of TIGR4, TIGR4 Δ*prp* and D39 were separated by SEC (Fig. 5B). Each fraction was then tested for cleavage of SUMO-L27. These results showed that fractions 8-9 in TIGR4 and D39, but not in TIGR4 Δ*prp*, were able to cleave SUMO-L27, whereas fractions 4-5 cleaved SUMO-L27 only in TIGR4 and TIGR4 Δ*prp*, but not in D39 (Fig. 5C). This result suggested that the unknown protease was present in TIGR4 fraction 4-5, while Prp was found in fractions 8-9, consistent with its known 12.8 kDa molecular weight.

Fractions 4-5 and 8-9 from TIGR4 wildtype, TIGR4 Δ*prp* and D39 were subsequently concentrated 10-fold and tested for cleavage of SUMO-L27 and SUMO-L27F12A (Fig. S4A). Efficient cleavage of SUMO-L27 by TIGR4 was seen in both fractions, whereas D39 only showed cleavage in fraction 8-9 (Fig. S4A). Cleavage of SUMO-L27 was greatly reduced in Δ*prp* and was only seen in fraction 4-5 (Fig. S4A). Cleavage of SUMO-L27F12A was observed in fraction 4-5 in TIGR4, albeit significantly reduced compared to SUMO-L27 (Fig. S4A). To confirm that the cleavage did not occur within the SUMO domain, we tested the unrelated SUMO fusion protein SUMO-gp44 (gp44 is a protein from *S. aureus* bacteriophage 80α) for cleavage by TIGR4, TIGR4 Δ*prp* and D39 lysates. No cleavage of SUMO-gp44 was observed, showing that the SUMO domain itself was not cleaved by any protease present in the lysates (Fig. S4B).

Fractions 4-5 from TIGR4, TIGR4 Δ*prp* and D39 were separated by SDS-PAGE, subjected to trypsin digestion, and analyzed by quantitative mass spectrometry (qMS) (Fig. 6A). We hypothesized that peptides corresponding to the unknown protease would be present in TIGR4 and in TIGR4 Δ*prp,* but absent from D39 (Fig. 6A). To identify potential candidates, we analyzed the qMS data using Scaffold Viewer, employing a 95% protein threshold, an 80% peptide threshold, and a minimum of two peptides to minimize nonspecific identifications. A total of 655 proteins were identified. Among these, 90 proteins were identified that had multiple peptides in both TIGR4 and TIGR4 Δ*prp*, but no detectable peptides in D39. Within this subset, only five proteins were annotated as known proteases or peptidases: ZmpA, ZmpB, ZmpC, X-pro dipeptidyl peptidase, and signal peptidase I. Since ZmpA, ZmpB, and ZmpC had already been tested and shown not to be responsible for L27/L27F12A cleavage (Fig. 5A), they were excluded. X-pro dipeptidyl peptidase and signal peptidase I were also excluded as candidates because the genes encoding both these proteins are present in the D39 genome. This analysis suggested that the protease responsible for cleavage of bL27 had not yet been identified and would be annotated as a “hypothetical protein” in the NCBI database. We therefore refined our search by excluding proteins with known functions, yielding 26 hypothetical proteins (Fig. 6A). BLAST analysis was performed for each of these proteins to identify any known or functionally similar homologs, which allowed us to exclude three additional candidates. Among the remaining 23 proteins, further analysis revealed that genes encoding 20 of those were present without any mutations or truncations in the D39 genome, making them less likely candidates (Fig. 6A). Thus, only three candidate proteins remained that were identified in TIGR4 and TIGR4 Δ*prp* but not in D39, and for which the genes were absent from the D39 genome: *SP_1793*, *SP_2093*, and *SP_1145* (Fig. 6A).

**Figure 6.**
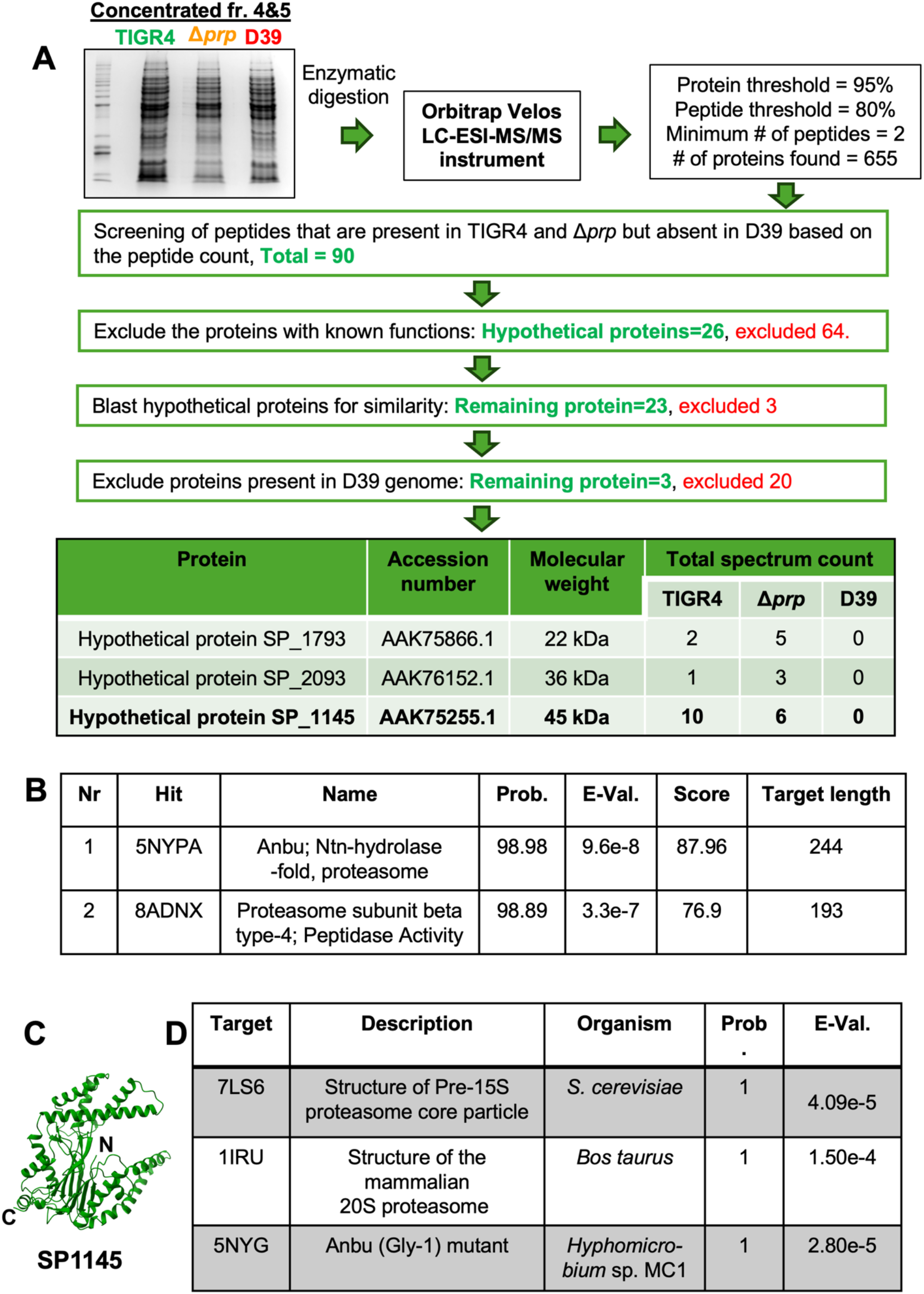
Identification of the alternate protease that cleaves bL27. **(A)** Flow chart of the qMS approach to identify the alternate protease, resulting in the three candidates SP1793, SP2093 and SP1145, showing the total spectrum count for each. **(B)** Top two hits from the HHPred server searched with the suspected protease SP1145 from the qMS analysis in (A). Probability, E-value, HHpred score and residue length of matched region are listed. **(C)** Ribbon diagram of the AlphaFold prediction model of SP1145. N- and C-termini are indicated. **(D)** Top three hits from the PDB database after FoldSeek search with the SP1145 Alphafold model. Probability and E-value are shown.

The protein sequences for *SP_1793*, *SP_2093* and *SP_1145* were subjected to structural analysis with HHpred [36] (Fig 6B), which revealed that the product of the *SP_1145* gene (protein SP1145), had homology with proteases related to eukaryotic 20S proteasome β subunits and to ancestral eubacterial homolog Anbu [30, 31], proteases that are responsible for degradation of misfolded proteins. No homology with predicted proteases was detected for the other two proteins. Furthermore, SP1145 was the only one of the three candidates that was identified by MS in purified ribosomes from TIGR4 and Δ*prp* (Table S2).

We generated a model for SP1145 in AlphaFold [37] (Fig 6C), which was submitted to the FoldSeek server [38]. This search also identified similarity to proteasome-like proteins from different species, including eukaryotic proteasome β subunits from *Saccharomyces cerevisiae* (PDB ID: 7LS6) and *Bos taurus* (PDB ID: 1IRU) and the bacterial Anbu protein from *Hyphomicrobium* sp. [31] (PDB ID: 5NYG; Fig 6D). Superposition of the SP_1145 model on these structures showed that while the core structure was conserved, SP_1145 had additional extended domains (Fig. S5A). Interestingly, FoldSeek also identified related structures predicted by AlphaFold in several bacteriophage genomes (Fig. S5B) suggesting that the *SP_1145* gene could have been acquired horizontally.

### 6. Deletion and complementation of *SP_1145* in *S. pneumoniae* TIGR4

The above results strongly implicated *SP_1145* as the gene encoding the alternate bL27 protease in TIGR4. To validate this result, we deleted *SP_1145* from TIGR4 using the easyJanus method (Fig. 7A, Fig. S3, Table 1). The resulting TIGR4 Δ*SP_1145* mutant showed a slight growth defect compared to the wildtype (Fig. 7B). The TIGR4 Δ*SP_1145* cell lysate was tested for cleavage of SUMO-L27 and SUMO-L27F12A, which showed that the Δ*SP_1145* strain was indeed incapable of cleaving SUMO-L27F12A, similar to D39 (Fig. 7C, Fig. 4C, D). In contrast, wildtype SUMO-L27 was cleaved normally in the Δ*SP_1145* mutant, presumably by Prp.

**Figure 7.**
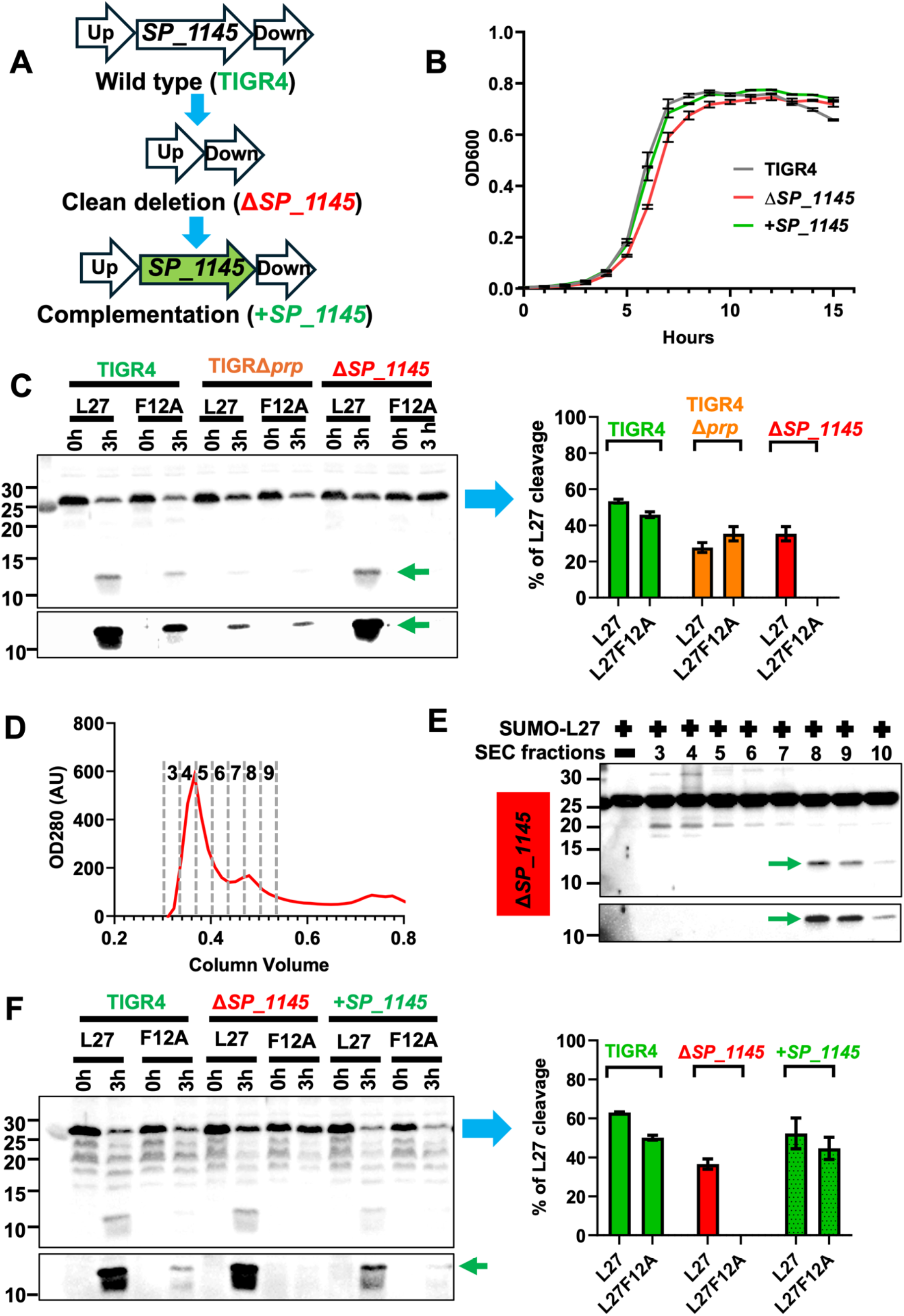
Deletion and complementation of *SP_1145*. **(A)** Flow chart of deletion and same-site complementation of *SP_1145* in the TIGR4 genome. **(B)** Growth curves for TIGR4 wildtype (gray), TIGR4 Δ*SP_1145* (red) and TIGR4 Δ*SP_1145* complemented with *SP_1145* in the same site (+*SP_1145*; green) (average of three measurements). **(C)** Cleavage of SUMO-L27 and SUMO-L27F12A by cell lysates from TIGR4, TIGR4 Δ*prp* and TIGR4 Δ*SP_1145*, separated by SDS-PAGE and probed with the anti-bL27 antibody. A higher exposure showing the bands corresponding to L27ΔN (green arrows) is shown below the main gel. Densitometric quantitation of cleavage is shown on the right. **(D)** SEC fractionation of the TIGR4 Δ*SP_1145* cell lysate. Fractions collected are indicated by the numbers. **(E)** Cleavage of SUMO-L27 by TIGR4 Δ*SP_1145* SEC fractions from (C), separated by SDS PAGE and probed with the anti-bL27 antibody. A higher exposure showing the L27ΔN band (green arrow) is shown below the main gel. **(F)** Cleavage of SUMO-L27 and SUMO-L27F12A by lysates of TIGR4 wildtype, TIGR4 Δ*SP_1145* and TIGR4 Δ*SP_1145* +*SP_1145*, separated by SDS-PAGE and probed with the anti-bL27 antibody. Higher exposure shown below; the cleaved L27ΔN is indicated by the green arrow. Densitometric quantitation of cleavage is shown on the right.

As described above, the Prp activity was found to reside in fractions 8-9 after SEC separation of a TIGR4 lysate, while the alternate protease was found in fractions 4-5 (Fig. 5C). We separated the Δ*SP_1145* mutant lysate by SEC (Fig. 7D), followed by testing the fractions in the cleavage assay (Fig. 7E). These results confirmed that deletion of *SP_1145* abolished activity in fractions 4-5, while activity was retained in fractions 8-9, similar to strain D39 (Fig. 5C). When the *SP_1145* gene was complemented back in the original locus, the ability to cleave SUMO-L27F12A was regained (Fig. 7F) and the growth recovered to wildtype levels (Fig. 7B). Collectively, these results strongly indicate that SP1145 is the alternate protease that cleaves bL27 in the TIGR4 Δ*prp* strain in the absence of Prp. We propose to name this new protease Rrp, for “<u>R</u>ibosome rescue <u>p</u>rotease.”

### 7. Cleavage of bL27 by Rrp in vitro

*SP_1145* was cloned from the TIGR4 genome into the pET21a vector and expressed in *E. coli* with a C-terminal His_6_ tag. When expressed in *E. coli* BL21(DE3) cells, the protein caused markedly slow bacterial growth, suggesting that the Rrp protein may be toxic to *E. coli*. Expression in BL21(DE3) pLysS cells resulted in a faint protein band upon IPTG induction, as detected by an anti-His_6_ antibody (Fig. 8A). The observed molecular weight of the detected protein was between 40 and 50 kDa, consistent with the expected Rrp mass of 44.8 kDa.

**Figure 8.**
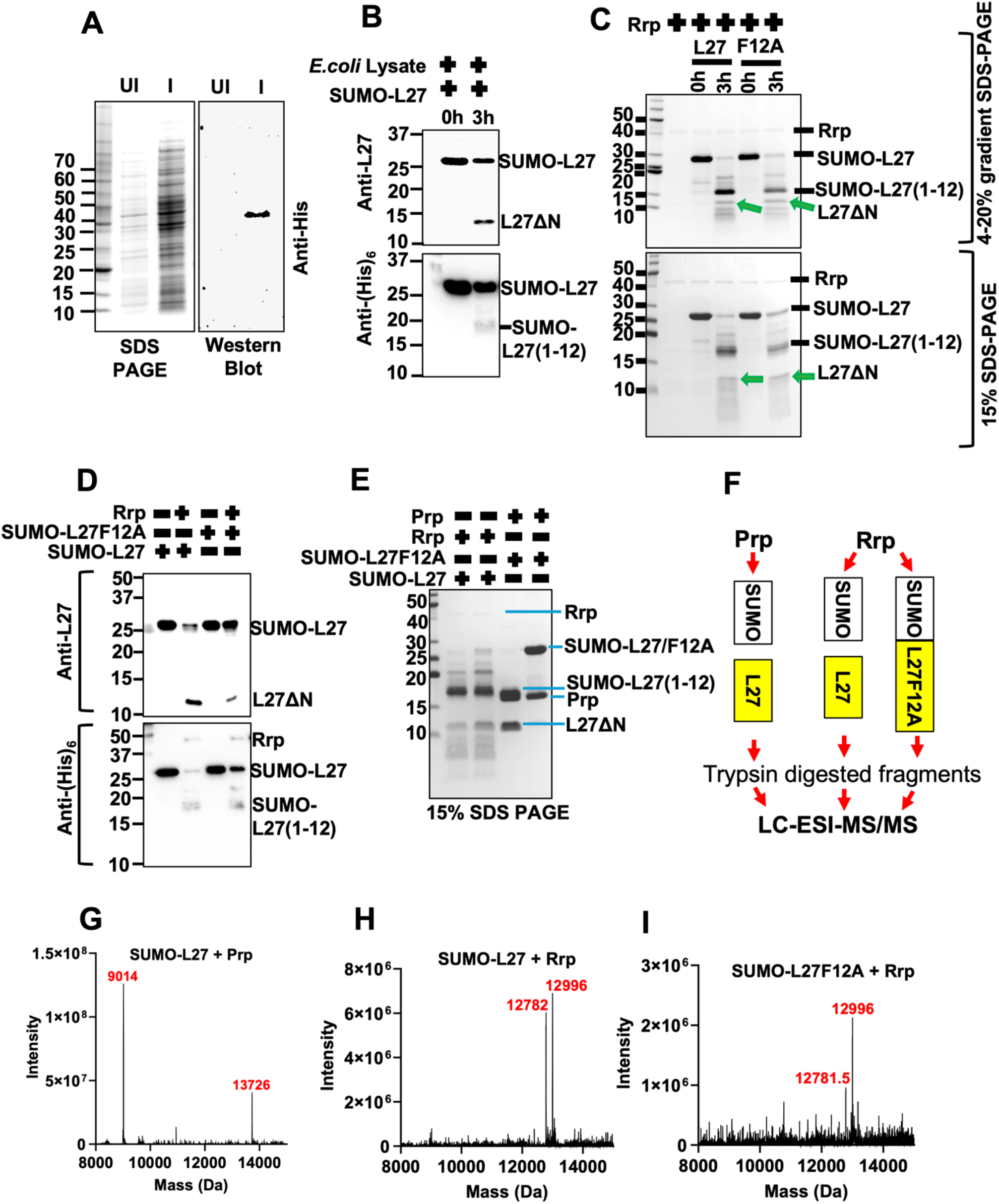
In vitro activity and cleavage analysis of Rrp. **(A)** Expression of His_6_-tagged Rrp (SP1145) in *E. coli*. Coomassie stained SDS-PAGE (left) of whole cell lysate before (Uninduced, UI) and after (Induced, I) IPTG induction. Western blot probed with anti-His_6_ antibody shown on the right. **(B)** Cleavage of SUMO-L27 by *E. coli* cell lysate (1 mg/ml) expressing Rrp, incubated for 3 h at 37 °C, separated by SDS-PAGE and probed by either anti-bL27 (top) or anti-His_6_ (bottom) antibody. The L27ΔN fragment detected by anti-bL27 antibody and the SUMO-L27(1-12) fragment detected by the anti-His_6_ antibody are indicated. **(C)** Purified Rrp was incubated with 3.4 µM SUMO-L27 or SUMO-L27F12A for either 0 h or 3 h at 37 °C and analyzed by 4-20% gradient (top) or 15% (bottom) SDS PAGE. Bands corresponding to Rrp, SUMO-L27 and SUMO-L27(1-12) are indicated. The bands corresponding to cleaved L27ΔN are marked by green arrows. **(D)** Cleavage of SUMO-L27 and SUMO-L27F12A by purified Rrp, incubated for 3 h at 37 °C, followed by SDS-PAGE separation and Western blotting with either anti-bL27 (top) or anti-His_6_ (bottom) antibodies. **(E)** Cleavage of SUMO-L27 and SUMO-L27F12A (10 µM) by either 10 µM purified Prp or 1 µM purified Rrp, incubated at 37 °C for 2 h, separated by 15% SDS-PAGE and Coomassie stained to provide samples for MS analysis. **(F)** Schematic representation of the samples from (E) that were analyzed by LC-ESI-MS/MS. **(G-I)** Mass spectra of peptides detected after cleavage of SUMO-L27 by Prp (G), cleavage of SUMO-L27 by Rrp (H) and cleavage of SUMO-L27F12A by Rrp (I). Masses of the most abundant peptides (Da) are indicated in red.

We then used the same lysate cleavage assay as above (Fig. 3A) to assess whether *E. coli* lysates expressing Rrp could cleave SUMO-L27. [We already confirmed that an *E. coli* lysate alone did not cleave SUMO-L27 in vitro (Fig. S2).] The reaction was separated by SDS-PAGE and probed by Western blot using the anti-bL27 and anti-His_6_ antibodies (Fig. 8B). The anti-bL27 antibody detected a band corresponding to L27ΔN, while an anti-His_6_ antibody detected a faint band corresponding to the SUMO-L27(1-12) fragment, concomitant with a reduction in intensity of the SUMO-L27 band (Fig. 8B). These results clearly showed that the Rrp-expressing lysate was capable of cleaving SUMO-L27.

We purified the His_6_-tagged Rrp protein by affinity chromatography. Although the yield was very low, a band corresponding to Rrp was visible by SDS-PAGE at the expected molecular weight (Fig 8C). The purified Rrp was incubated with either SUMO-L27 or SUMO-L27F12A for 3h at 37 °C, followed by SDS-PAGE separation (Fig. 8C). This experiment showed that Rrp was capable of cleaving both SUMO-L27 and SUMO-L27F12A; however, cleavage of the L27F12A mutant was less efficient, consistent with previous results (Fig. 4). Furthermore, several bands were present, suggesting that the cleavage was not very precise. Western blot analysis of the same samples using anti-bL27 or anti-His_6_ antibodies further confirmed this difference, demonstrating that the L27F12A mutant can be cleaved by Rrp, but less efficiently than wildtype bL27 (Fig. 8D). The anti-His_6_ Western blot also confirmed the presence of the His_6_-tagged Rrp in the reaction mixture (Fig. 8D). A direct comparison with Prp cleavage highlights the differences between the two proteases: Prp cleaves SUMO-L27 very precisely, but is incapable of cleaving SUMO-L27F12A, whereas Rrp cleaves both substrates, with a slight preference for the wildtype SUMO-L27 protein (Fig. 8E).

The reaction mixtures from the cleavage of SUMO-L27 and SUMO-L27F12A with the purified Rrp protein were subjected to full-length MS to determine the cleavage site(s) (Fig. 8F). As a control, SUMO-L27 was cleaved with purified Prp. Cleavage of SUMO-L27 by purified Prp resulted in two major peaks in the mass spectrum at the expected masses for L27ΔN (9,014 Da) and SUMO-L27(1-12) (13,726 Da) (Fig. 8G, Fig. S6A, B). Cleavage by Rrp was much less specific; however, two major peaks were observed in the mass spectra at 12,782 and 12,996 Da for both SUMO-L27 and SUMO-L27F12A, corresponding to SUMO-L27 cleaved after Met 4 and Leu 6 of bL27, respectively (Fig. 8H-I; Table S3). When the region from 8,500-9,500 Da in the mass spectrum was expanded, peptides corresponding to the native F|A cleavage site as well as several other bL27-derived peptides were observed (Fig. S7-S8), but did not rise significantly above the background, again indicating that Rrp exhibited relaxed cleavage specificity.

## DISCUSSION

Ribosomal protein bL27 is a key component of the eubacterial ribosome, and consequently *rpmA* is an essential gene in *S. pneumoniae* and other Gram-positive bacteria [39–43]. We previously showed that ribosomal protein bL27 in Gram-positive organisms has an N-terminal extension that is cleaved by the Prp protease [18]. Using a dual expression system where the wildtype *rpmA* gene could be turned on and off, Wall et al [22] showed that *S. aureus* cells expressing either the N-terminally truncated, mature form of bL27 or the uncleavable mutant L27F12A were non-viable in the absence of expression of the wildtype allele, showing that both the presence of the extension and its cleavage were essential events. Indeed, the location of the bL27 N-terminus in the peptidyl transferase center is consistent with a strong effect on translation if cleavage is blocked [19, 44].

Since Prp was shown to be the protease responsible for cleaving bL27 [18], the *prp* gene was likewise assumed to be essential. Indeed, overexpression of protease-inactive mutant Prp from a plasmid was detrimental to cell growth in *S. aureus*, even in the presence of the wildtype *prp* allele [18]. This strain also displayed a distinct ribosome profile in which 40S ribosomal particles accumulated, suggesting that bL27 cleavage by Prp controlled some step in the ribosome assembly pathway. In light of these results, we were surprised to find that the *S. pneumoniae* TIGR4 Δ*prp* mutant was viable and displayed no ribosomal assembly defect (Fig. 2B, C). Moreover, ribosomes formed in the Δ*prp* mutant contained cleaved bL27 (Fig. 2F, G), suggesting that another protease was able to carry out this cleavage in the absence of Prp.

In this study, we identified this alternate protease as the product of gene *SP_1145*, encoding a protein of hitherto unknown function, that we have named Rrp, for <u>R</u>ibosome rescue <u>p</u>rotease. Our analysis showed that Rrp belongs to the Ntn hydrolase family of proteases and is related to eukaryotic proteasome β subunit proteases and their ancestral eubacterial equivalent Anbu [30, 31]. Anbu-like proteins have not been described in streptococci, which like other Firmicutes were assumed to lack the *anbu* gene [30]. These proteins are assumed to serve to degrade misfolded proteins in a process that is likely coupled to translation. Unlike Prp, Rrp was also able to cleave the L27F12A mutant protein, albeit less efficiently than bL27, indicating that it is a less specific process than cleavage by Prp. Indeed, we could not identify a unique cleavage site for Rrp on bL27 by MS. These results suggest that Rrp, like Anbu, is a general purpose protease that is capable of detecting and removing aberrant proteins, including the uncleaved N-terminal tail of bL27 present on the ribosome.

We have shown that *S. pneumoniae* TIGR4 possesses at least one protease (Rrp) in addition to Prp that can cleave the N-terminus of bL27, whereas *S. pneumoniae* D39 or *S. aureus* only have Prp (Fig. 9). In TIGR4 in the absence of Prp, the Rrp protein rescues the ribosomes by cleaving bL27 (Fig. 9). The *SP_1145* gene is present in TIGR4, but not in D39 (Fig. 9). Comparison of the TIGR4 and D39 genomes suggests that *SP_1145* is associated with a phage-like mobile genetic element (Fig. 10A, Table S4) and may have been acquired horizontally (Fig. 10B). Although *SP_1145* homologs are found in various streptococci, the element itself is not particularly common: out of the 18 *S. pneumoniae* sequences in the PneumoBrowse server (https://pneumobrowse2.veeninglab.com), only PJ755/1 has a mobile genetic element in the same locus, and TIGR4 is the only one that has the *SP_1145* gene.

**Figure 9.**
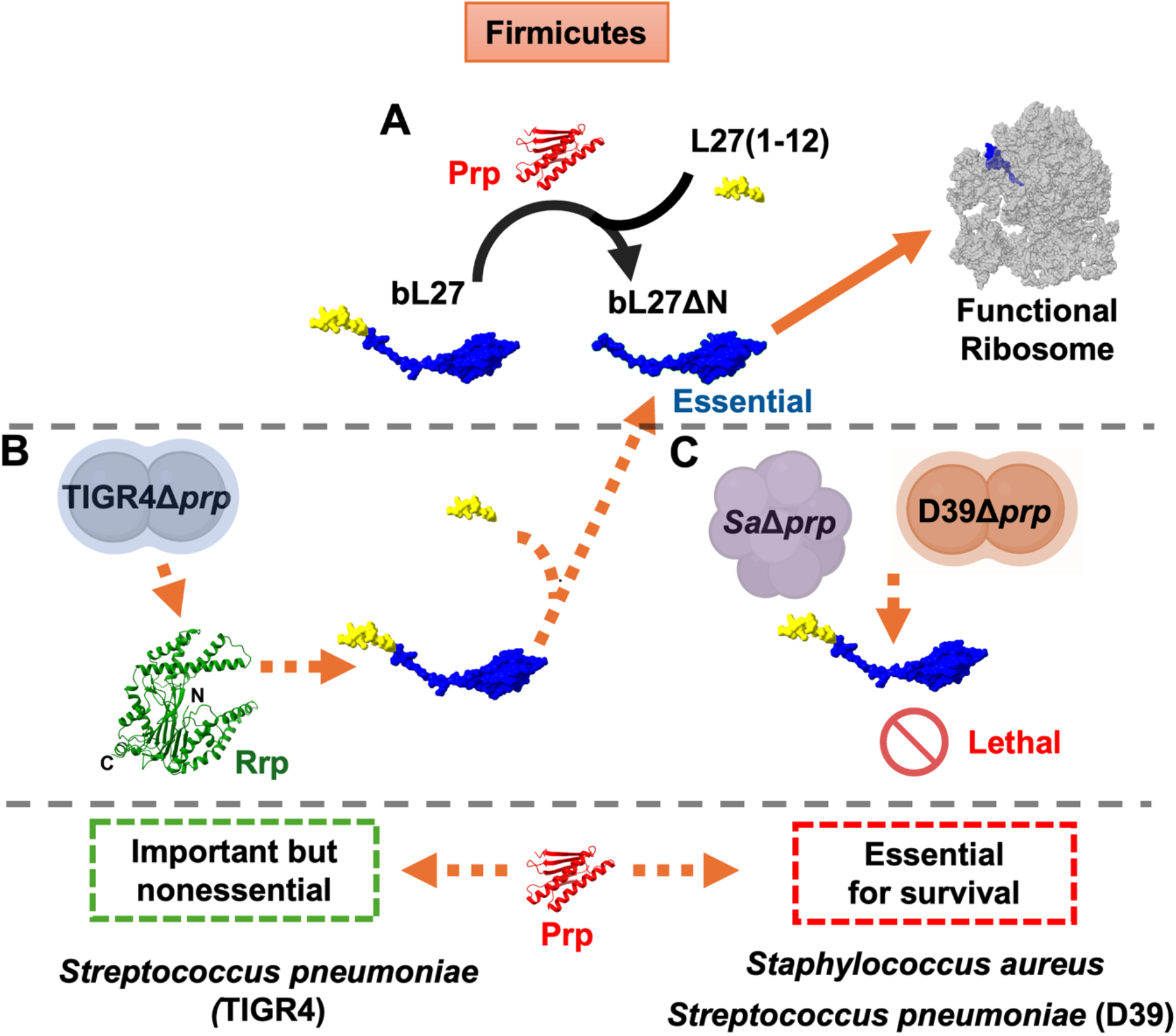
Model for bL27 cleavage in Firmicutes. **(A)** bL27 is normally cleaved by Prp, and only cleaved bL27 (L27ΔN) is incorporated into functional ribosomes. **(B)** In *S. pneumoniae* strain TIGR4, Rrp protease can cleave bL27 in the absence of Prp in a Δ*prp* deletion strain (TIGRΔ*prp*). **(C)** In *S. pneumoniae* strain D39 and in *S. aureus*, the *prp* deletion (Δ*prp*) is deleterious.

**Figure 10.**
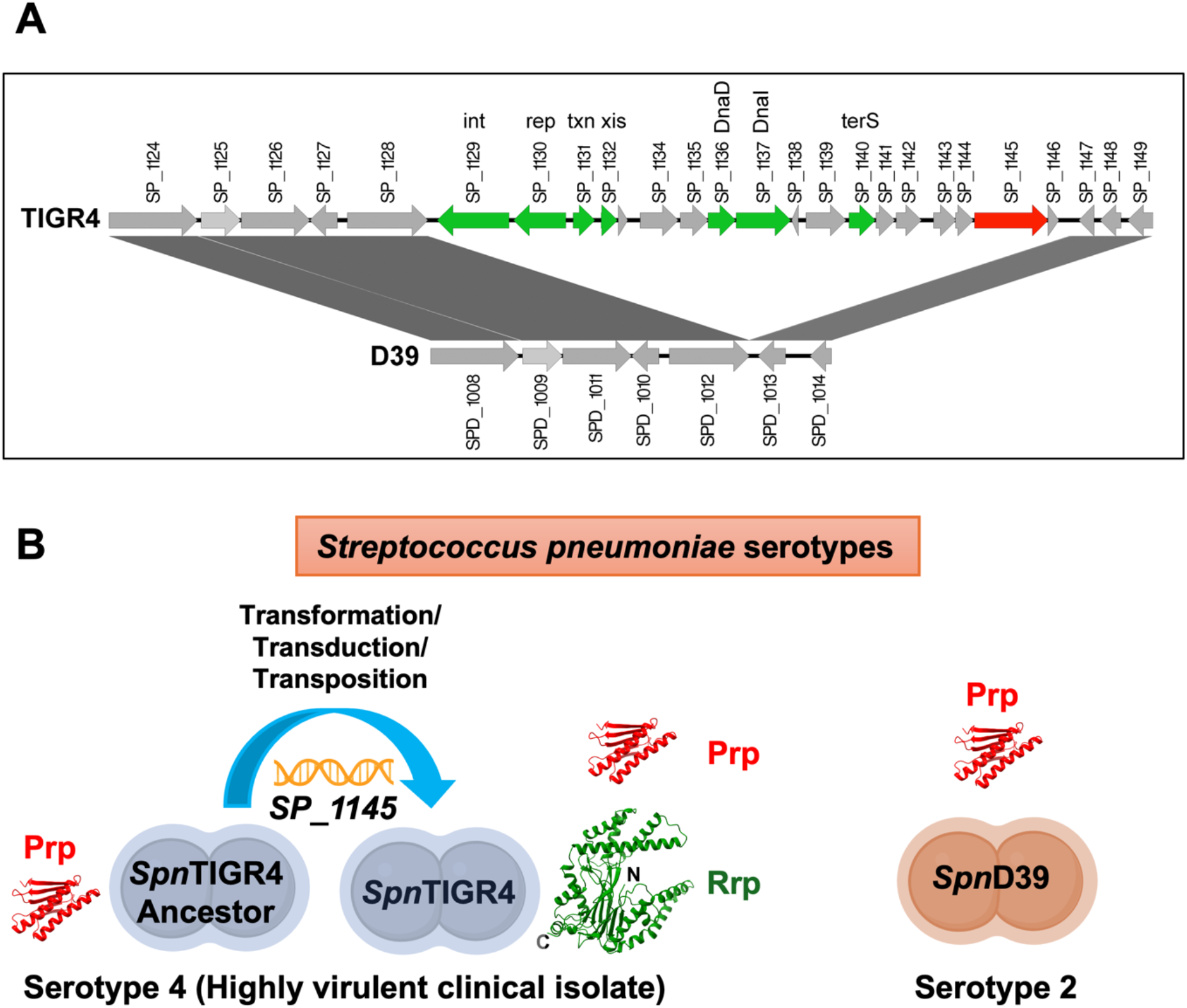
Genomic region around *SP_1145*. **(A)** Schematic diagram showing that *SP_1145* (red arrow) is located in a gene cluster in TIGR4 that is absent from D39. Phage-related genes detected by HHpred are indicated by green arrows. These genes include (top PDB hits from HHpred in parentheses): *int*, integrase (PDB ID: 6EMY); *rep*, repressor (7ZCV); *txn*, transcriptional regulator (7N1N); *xis*, excisionase (8DGL); *dnaD* (8OJJ) and *dnaI* (4M4W), parts of the primer complex; *terS*, small terminase (7JOQ). Figure generated with EasyFig [50]. **(B)** Model for horizontal acquisition of *SP_1145* by TIGR4 (serotype 4). Strain D39 (serotype 2) only has *prp*.

The similarity of the Rrp AlphaFold model to numerous predicted proteins from bacteriophages identified by the FoldSeek server also suggests that the gene may have originated in phages. Since Rrp is absent from strain D39, deletion of *prp* is lethal, and we were unable to recover viable Δ*prp* colonies by recombination (Fig. 4F). Indeed, the *prp* gene (*SPD_0990*) was found to be essential in D39 in a CRISPRi screen [39]. *S. aureus*, where *prp* is also essential, likewise lack an *SP_1145* homolog. Recently, Treerat et al [45] described a mutation in the *S. mutans prp* gene (*SMU_848*) that affected biofilm formation and glucan utilization. Although they did not characterize the effect of this mutant on bL27 cleavage or ribosome assembly, the lack of an effect on growth suggests that *S. mutans* also encodes another protease capable of cleaving bL27 [45]. *S. mutans* does not encode an Rrp homolog, so presumably the cleavage of bL27 in the absence of Prp in this species is done by some other protease. Overall, our results point to the acquisition of compensatory mechanisms capable of carrying out an essential processing step in the ribosome assembly pathway in the absence of Prp.

Why these bacteria have the N-terminal extension on bL27 in the first place in still unclear. It is possible that it serves a role in regulation of ribosomal activity, e.g. under conditions of starvation or other stressors, perhaps offering protection against antibiotics that target the ribosome. The presence of the extension in some staphylococcal bacteriophages [18] is especially intriguing, suggesting that the sequence is related to an ancient defense mechanism against phages or other environmental stressors.

Prp has been proposed as a target for novel antibiotics specific to Gram-positive bacteria [18, 22, 28, 29]. While this approach may have merit in some species, such as *S. aureus*, it is clear that some bacterial pathogens, including some strains of *S. pneumoniae*, have alternate pathways for carrying out the cleavage of bL27, rendering this approach less effective. Moreover, Prp-suppressing drugs could potentially promote the development of resistance via horizontal dissemination of Rrp or other proteases. These aspects need to be considered before Prp can be considered a therapeutic target.

## EXPERIMENTAL PROCEDURES

### 1. Cloning and site-directed mutagenesis in *E. coli*

*S. pneumoniae* strain TIGR4 (GenBank ID: AE005672)[46] gene *prp* (*SP_1106*) was cloned into the pET21a vector using In-Fusion cloning (Takara), incorporating an N-terminal T7 leader sequence followed by the *prp* coding region, a myc tag, a thrombin cleavage site, and a C-terminal hexahistidine (His_6_) tag (Table 1). Similarly, the *S. pneumoniae* TIGR4 gene *rpmA* (*SP_1107*) was cloned into the pE-SUMO vector, which encodes an N-terminal His_6_ tag and a SUMO domain, also using In-Fusion cloning. Site-directed mutagenesis of *rpmA* in pSUMO-L27 was performed using the In-Fusion cloning method (Takara). The *SP_1145* gene was cloned by the same method as *prp* except the N-terminal T7 tag and the TTG start codon of *SP_1145* was replaced by an ATG start codon, and only a His_6_ tag was included at the C-terminus.

### 2. Protein expression and purification

His_6_-tagged recombinant proteins were expressed in *E. coli* BL21(DE3) or *E. coli* BL21(DE3) pLysS cells following induction with 1 mM IPTG. The expressed proteins were purified using HisPur™ Cobalt Resin (Thermo Scientific). Affinity-purified proteins were concentrated using Amicon centrifugal filter units and further purified by size-exclusion chromatography using Superdex™ 75 Increase columns in a Bio-Rad NGC chromatography system. Eluted protein fractions were collected and concentrated using Amicon filters.

### 3. Generation of *S. pneumoniae* mutants

A 2,873 bp kanamycin resistance cassette was constructed by assembling a 1,010 bp left homology arm (upstream of *prp*), a 1068 bp right homology arm (downstream of *prp*), and a 795 bp kanamycin resistance gene (*aphIII)*. These components were joined via In-Fusion PCR and cloned into the pPEPZ-Plac plasmid [47]. The recombinant plasmid was transformed into *S. pneumoniae* TIGR4, and transformants were selected on blood agar plates containing kanamycin, yielding the TIGR4 Δ*prp* strain MN001 (Table 1).

Deletions of candidate protease genes (Table 1) were produced in TIGR4s using the Janus cassette [34] as previously described [48] or by the easyJanus method [35]. For the latter approach, synthetic DNA fragments were obtained from Twist Bioscience (South San Francisco, CA, USA). DNA assembly was performed using the NEBuilder HiFi DNA Assembly Master Mix kit (New England Biolabs). The assembled constructs were introduced into competent cells in a single transformation step, and transformants were selected on kanamycin-containing agar plates, as previously described [35]. A *SP_1145* complementation strain in which the *SP_1145* gene was put back into the original locus was generated using the same method. All deletion and complementation strains were confirmed by whole-genome sequencing.

### 4. Bacterial growth assay

Growth assays for wild-type and mutant *S. pneumoniae* strains were performed in Todd-Hewitt broth supplemented with 2% yeast extract (THY). Briefly, cells were grown overnight on blood agar plates (Remel) at 37 °C with 5% CO₂. One-fourth of the plate surface was scraped using a sterile cell scraper and used to inoculate 10 mL of THY as a starter culture. The starter culture was grown until it reached an optical density (OD) of 0.3 at 600 nm wavelength (A_600_). A 2.5% inoculum from the starter culture was used to inoculate 200 mL of fresh THY medium. Bacterial growth was monitored by measuring A_600_ every hour for 10 h. All growth curves were performed in triplicate.

### 5. Ribosome purification

For ribosome purification, 500 mL cultures of *S. pneumoniae* TIGR4 and TIGR4 Δ*prp* were grown to A_600_ = 0.6–0.7 OD and harvested by centrifugation. Cell pellets were resuspended in 15 mL lysis buffer (20 mM Tris-HCl, pH 7.5; 50 mM NH_4_Cl; 15 mM MgCl₂; 0.5 mM EDTA; 6 mM β-mercaptoethanol) and lysed by three passes through an Avestin Emulsiflex B15 homogenizer. The lysate was clarified by centrifugation at 25,000 × g for 30 min at 4 °C. The clarified lysate (12 mL) was layered onto a 12 mL sucrose cushion (1.1 M sucrose in lysis buffer) and centrifuged at 50,000 rpm for 16 h at 4 °C in a Ti70 rotor. After centrifugation, the supernatant was discarded, and the ribosomal pellet was resuspended in 10 mL of lysis buffer. A 150 µL aliquot of the ribosome preparation was then layered onto a 5–40% (w/v) sucrose gradient prepared in lysis buffer and centrifuged at 75,500 × g for 16 h at 4 °C in an SW41 rotor. Fractions containing 70S ribosomal particles were identified, pooled, dialyzed against lysis buffer, and concentrated using a 100 kDa molecular weight cutoff membrane. The ribosome samples were negatively stained with 1% (w/v) uranyl acetate and examined by electron microscopy using an FEI Tecnai F20 transmission electron microscope.

### 6. Mass spectrometry

Purified SUMO-L27 or SUMO-L27F12A was mixed with either purified Prp or semi-purified Rrp (SP1145) and incubated at 37 °C for 1 h to allow cleavage. The resulting cleavage products were analyzed by intact mass spectrometry. Samples were desalted using a Waters Acquity BEH 200 Å 1.7um, 2.1 × 150 mm column with 0.1% formic acid as the mobile phase, at a flow rate of 100 µL/min. Data were acquired on a Waters Synapt G2-S(i) mass spectrometer operated in positive ion, resolution mode. Raw m/z spectra were deconvoluted to neutral mass using the MaxEnt 1 module within Waters MassLynx software. To examine the state of bL27 in ribosomes, TIGR4 wildtype and Δ*prp* 70S ribosomes were separated by SDS-PAGE on a 10% Bis-Tris gel and stained with colloidal Coomassie blue. A segment corresponding to the visible bands was extracted, digested with trypsin and analyzed by LC-ESI-MS/MS using a Thermo Orbitrap Velos Pro spectrometer. Data was quantitated as the total spectrum count, using a protein threshold of 95%, a peptide threshold of 80%, and a minimum number of peptides 2, and analyzed using the Scaffold Viewer. Full cell lysates of TIGR4, TIGR4 Δ*prp* (MN001) and D39 were analyzed similarly, by separating the proteins by SDS-PAGE, extracting the whole lane in three segments and digesting with trypsin, followed by MS analysis.

### 7. In vitro cleavage assay

Purified Prp or Rrp was mixed with equimolar amounts of purified SUMO-L27 or SUMO-L27F12A in assay buffer (25 mM HEPES pH 7.4, 0.3 M NaCl) and incubated at 37 °C for 1 h before boiling in Laemmli sample buffer. A control sample (0 h) was boiled in sample buffer immediately after mixing. Samples were separated by 15 % SDS-PAGE. For cleavage by cell lysates, the cells were harvested at A_600_=0.6 OD, resuspended in assay buffer and lysed in an Avestin Emulsiflex B15 homogenizer. The lysates were clarified by centrifugation, and the supernatants were filtered through a 0.2 µm membrane filter. Protein concentration in the lysate was determined using the Bradford assay (Bio-Rad) and normalized to 2 mg/mL. For the cleavage assay, 3.5 µM of purified SUMO-L27 or SUMO-L27F12A in assay buffer was mixed with the cell lysate and incubated at 37 °C for varying time intervals with gentle rotation. The control sample (0 h) was boiled instantly with Laemmli loading buffer after mixing. Samples were separated by 15% SDS-PAGE and analyzed by immunoblotting using either an anti-His_6_ primary antibody (Invitrogen, 37-2900) or a custom rabbit anti-*S. aureus* bL27 antibody [18]. A horseradish peroxidase (HRP)-conjugated anti-rabbit secondary antibody was used for detection. Signal development was carried out using a chemiluminescent substrate composed of luminol and hydrogen peroxide (Fisher Scientific). The amount of digestion was quantified from the blots by densitometric scan using ImageJ by comparing the intensity of the band corresponding to uncleaved protein after 3 h to the band at 0 h according to the following formula:

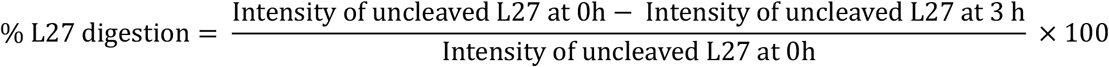

## DATA AVAILABILITY

All data are contained within the manuscript. Additional information and requests for reagents and resources are available from the corresponding author.

## SUPPORTING INFORMATION

This article contains supporting information including additional gel images and mass spectrometry data.

## ACKNOWLEDGEMENTS

We are grateful to Dr. Jessica Scoffield at UAB for providing strains and plasmids, and for assistance with genetics and bacteriology methods. We appreciate the assistance of Drs. Kyoko Kojima and Jim Mobley in the UAB Proteomics core for mass spectrometry services, Dr. Peter Prevelige for help with full-length MS and Dr. James Kizziah in the UAB Cryo-EM Facility (CEMF) for assistance with electron microscopy.

## FUNDING AND ADDITIONAL INFORMATION

CEMF is supported by the Institutional Research Core Program, The National Institutes of Health (NIH) grant S10 OD024978 to T.D. and the O’Neal Comprehensive Cancer Center (NIH grant P30 CA013148). This work was supported by an AMC21 pilot grant from the UAB Heersink School of Medicine and NIH grant R21 AI187107 to T.D., and by NIH grant R01 AI114800 to C.J.O. The content is solely the responsibility of the authors and does not necessarily represent the official views of the National Institutes of Health.

## AUTHOR CONTRIBUTIONS

**Amarshi Mukherjee:** Methodology, Investigation, Visualization, Writing—Original Draft, Writing— Review & Editing. **Mohamed O. Nasef:** Methodology, Investigation. **Patrick M. Lindstrom:** Investigation. **Nehaal Akavaram:** Investigation. **Eriel Martinez:** Investigation. **Vipin Chembilikandy:** Methodology, Investigation. **Carlos J. Orihuela:** Supervision, Funding Acquisition. **Terje Dokland:** Conceptualization, Supervision, Funding Acquisition, Writing—Original Draft, Writing—Review & Editing.

