## Supplemental Figures and Tables for "Rescue of ribosomal protein bL27 in *Streptococcus pneumoniae* TIGR4 by an alternate protease"

**Contains:**

**SUPPLEMENTARY FIGURES S1–S8**
**SUPPLEMENTARY TABLES S1–S4**
**SUPPLEMENTARY REFERENCES**

**Figure S1:**

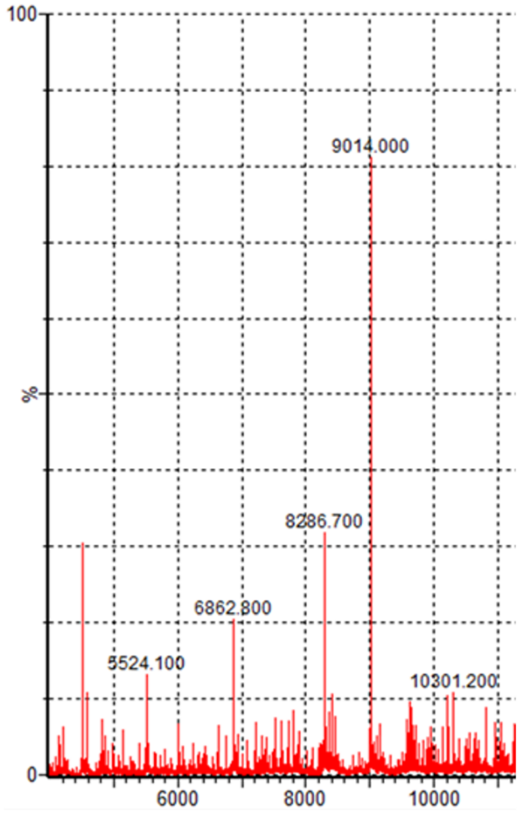


**Figure S1. Mass spectrometry of cleaved SUMO-L27.** Intact protein analysis by electrospray ionization mass spectrometry (ESI-MS) of SUMO-L27 cleaved with Prp, showing a peak at 9,014 Da, corresponding to the expected mass of bL27 cleaved between Phe 12 and Ala 13 (predicted mass 9,014.15 Da).

#### Figure S2:

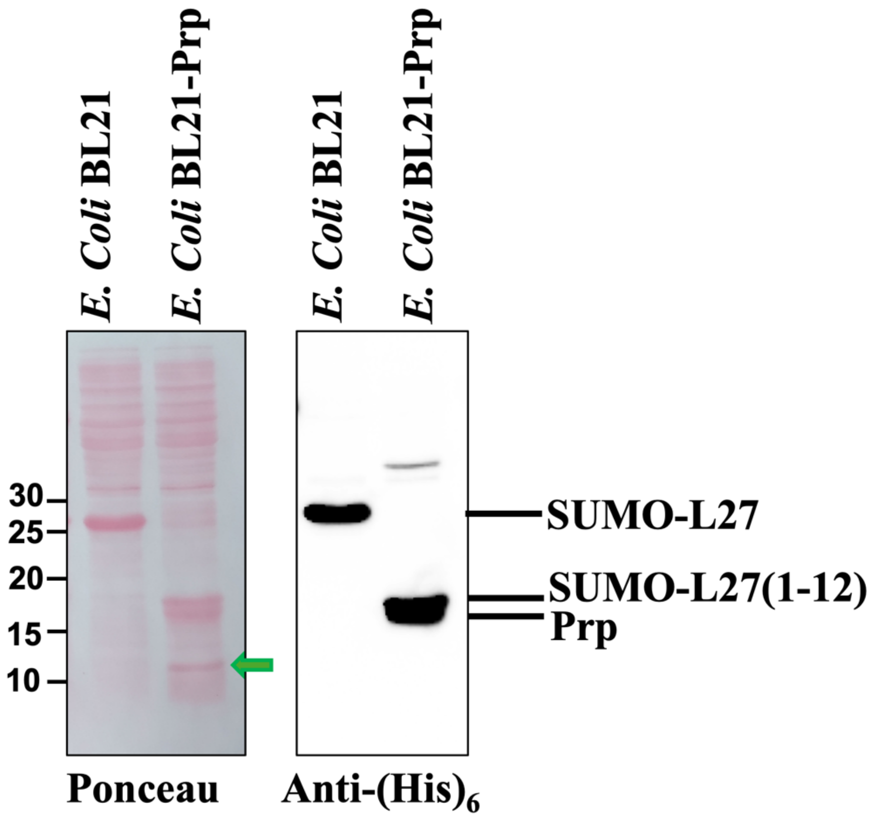


**Figure S2. Cleavage of SUMO-L27 by *E. coli* cell lysates.** SDS-PAGE separation of SUMO-L27 after incubation with a lysate of *E. coli* BL21(DE3) or *E. coli* BL21(DE3) overexpressing Prp from the pMN009 plasmid (BL21-Prp). The left panel shows the Ponceau stained membrane; the right panel shows a Western blot probed with the anti-$His_6$ antibody. Bands corresponding to Prp, the uncleaved SUMO-L27, and the N-terminal SUMO-L27(1-12) cleavage product are indicated on the Western blot. The cleaved L27ΔN fragment is visible by Ponceau stain (green arrow).

#### Figure S3:

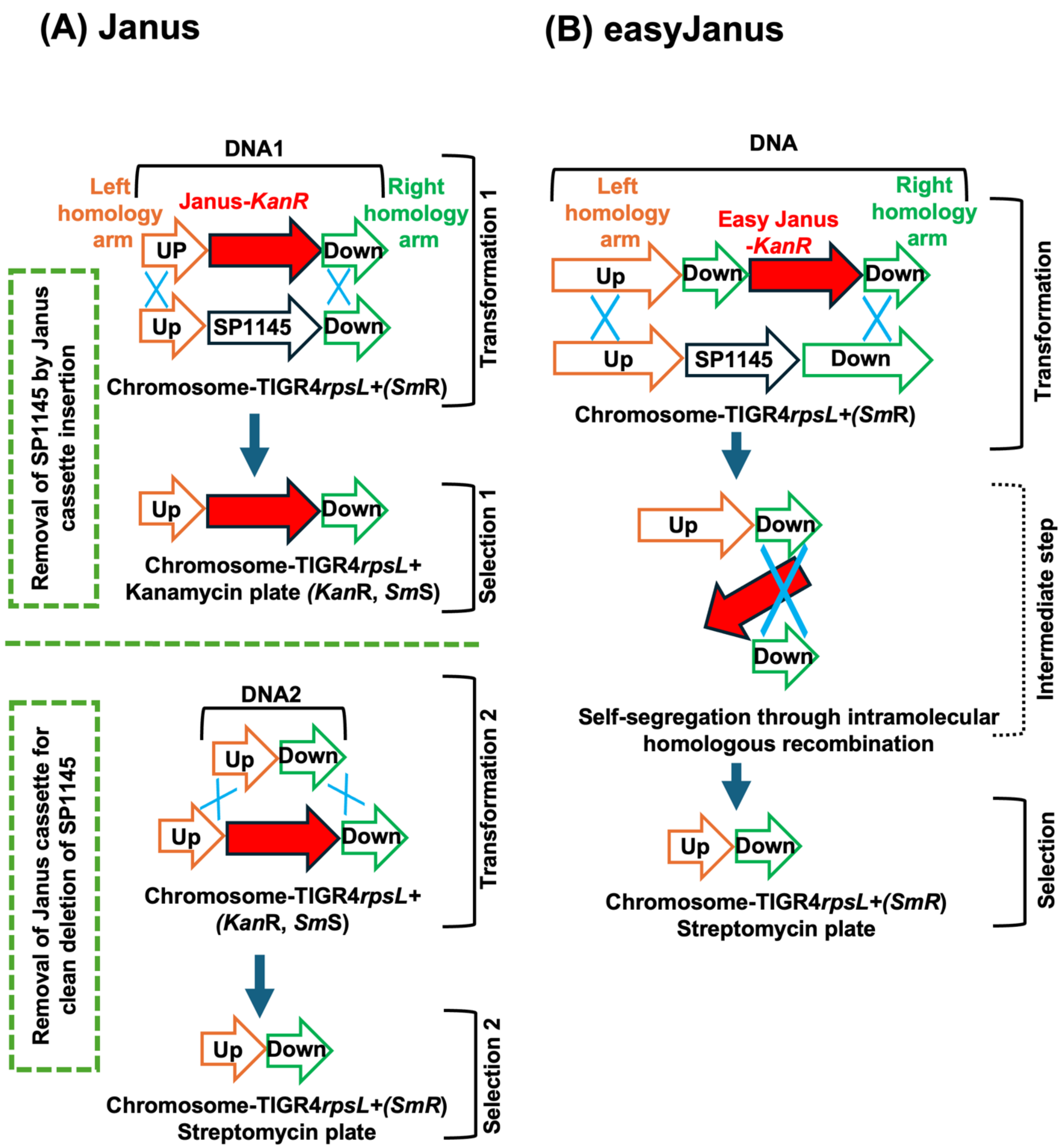


**Figure S3. Genetic recombination with Janus and easyJanus.** Schematic representation of gene replacement using the Janus cassette in the TIGR4s background, in which the *rpsL*+ mutation confers streptomycin resistance [1]. The Janus cassette (red color) confers kanamycin resistance and streptomycin sensitivity when it is inserted through homologous recombination into the gene locus of interest. **(A)** In the traditional Janus method [1], a second transformation is used to excise the Janus cassette, regaining kanamycin sensitivity and streptomycin resistance that is selected by plating on streptomycin, resulting in a clean deletion with no antibiotic marker. **(B)** In the easyJanus method [2], self-recombination leads to excision of the Janus cassette in a single step.

### Figure S4:

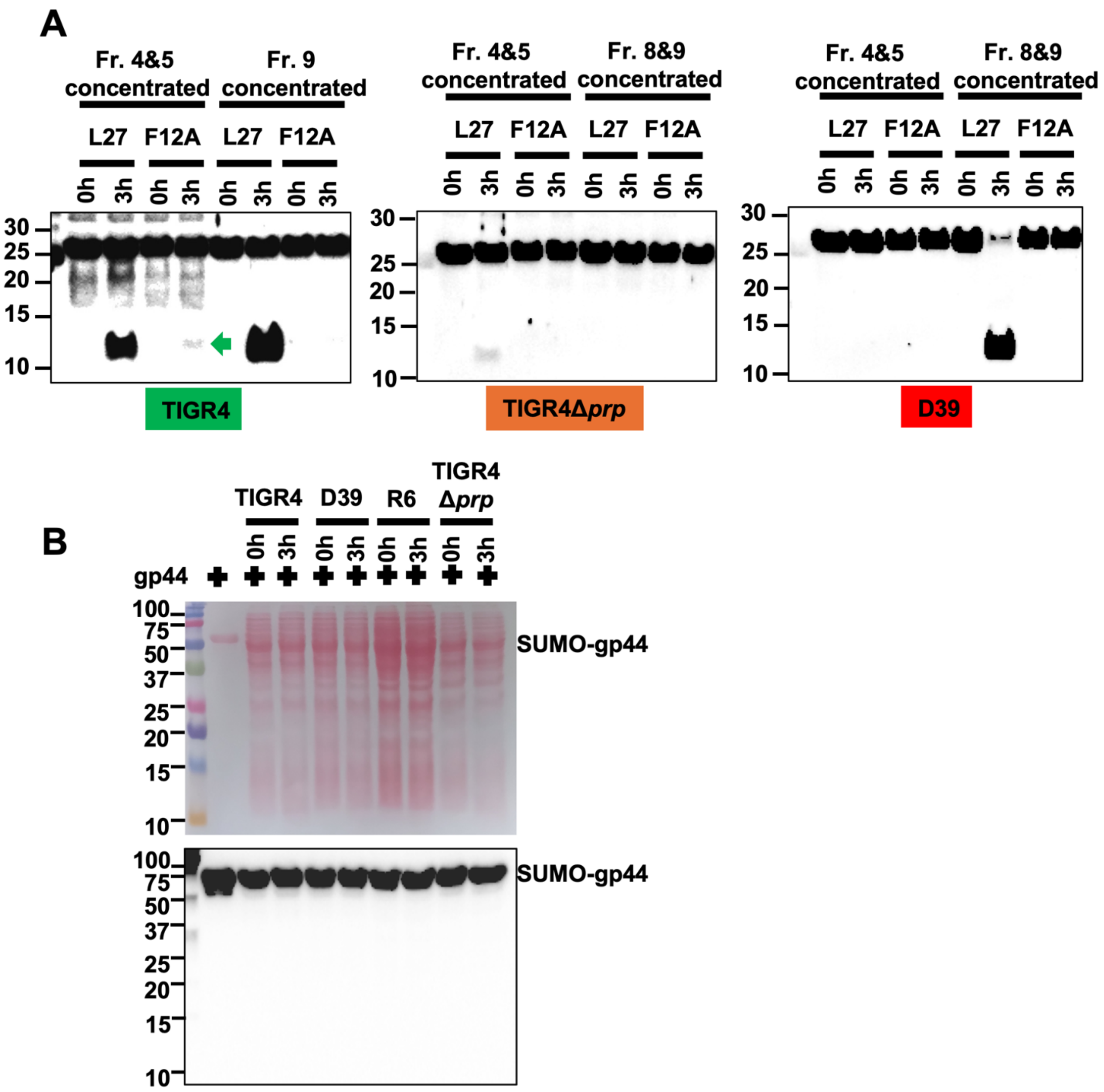


**Figure S4. Cleavage of SUMO-L27 and SUMO-L27F12A**. **(A)** Cleavage of SUMO-L27 and SUMO-L27F12A by 10-fold concentrated SEC fractions 4-5 and (8-)9 from TIGR4 wildtype, TIGR4 Δ*prp* and D39 cell lysates, separated by SDS-PAGE and probed by anti-bL27 antibody. The C-terminal L27ΔN fragment cleaved from SUMO-L27F12A by TIGR4 fraction 4-5 is indicated by the green arrow. **(B)** Control for cleavage, in which SUMO-gp44—a SUMO fusion with an unrelated *S. aureus* phage 80α protein—was incubated with *S. pneumoniae* strain TIGR4, D39, R6 or TIGR4 Δ*prp* lysates for 3 h, separated by SDS-PAGE and blotted with anti-bL27 antibody. The Ponceau stained membrane is shown above the blot. No cleavage of SUMO-gp44 could be detected.

### Figure S5:

**A**

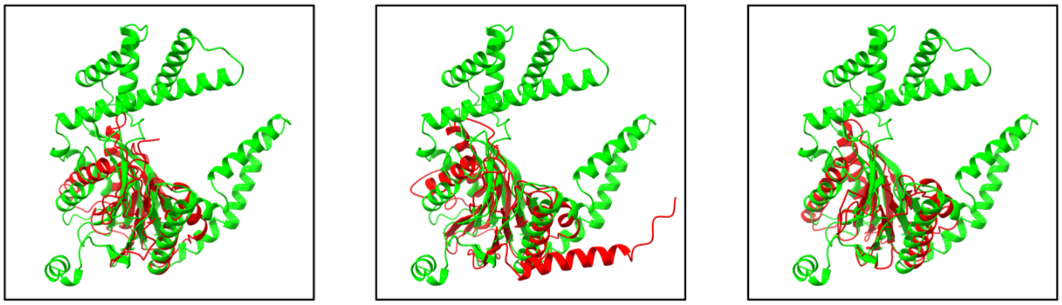

1IRU_Rrp 5NYG_Rrp 7LS6_Rrp

**B**

| Target | Description | Organism | Prob. | E-Val. |
|---|---|---|---|---|
| A0A8S5LJV3 | Proteasome beta type-2 subunit | Siphoviridae sp. Ct3o911 | 1 | 9.75e-7 |
| A0A142KAQ6 | Uncharacterized protein | Nymphadoravirus | 1 | 5.56e-4 |
| A0A0B5H8V9 | Proteasome endopeptidase complex | Jesfedecavirus | 1 | 2.99e-3 |

**C**

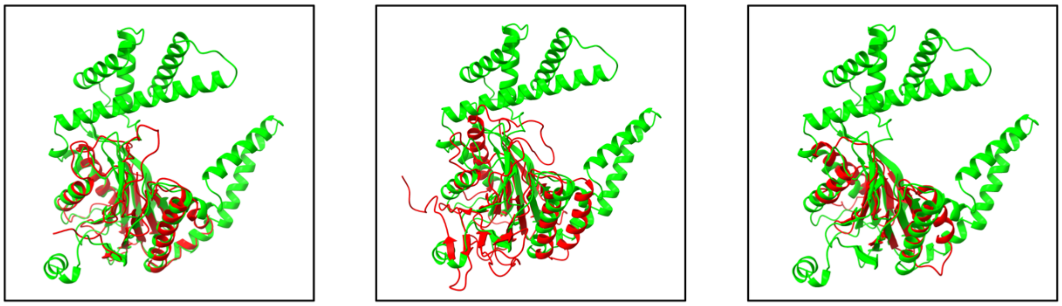

A0A8S5LJV3_Rrp A0A142KAQ6_Rrp A0A0B5H8V9_Rrp

**Figure S5. Structural homologs of SP1145. (A)** Ribbon diagrams of the SP1145 AlphaFold model (green) superimposed on proteins 1IRU, 5NYG and 7LS6 (red) identified by FoldSeek. **(B)** List of predicted bacteriophage proteins with high similarity to SP1145 identified in the Big Fantastic Virus Database (BFVD) by FoldSeek. The probability and E-value are listed. **(C)** Ribbon diagrams of the SP1145 AlphaFold model (green) superimposed on the predicted protein structures from (B).

### Figure S6:

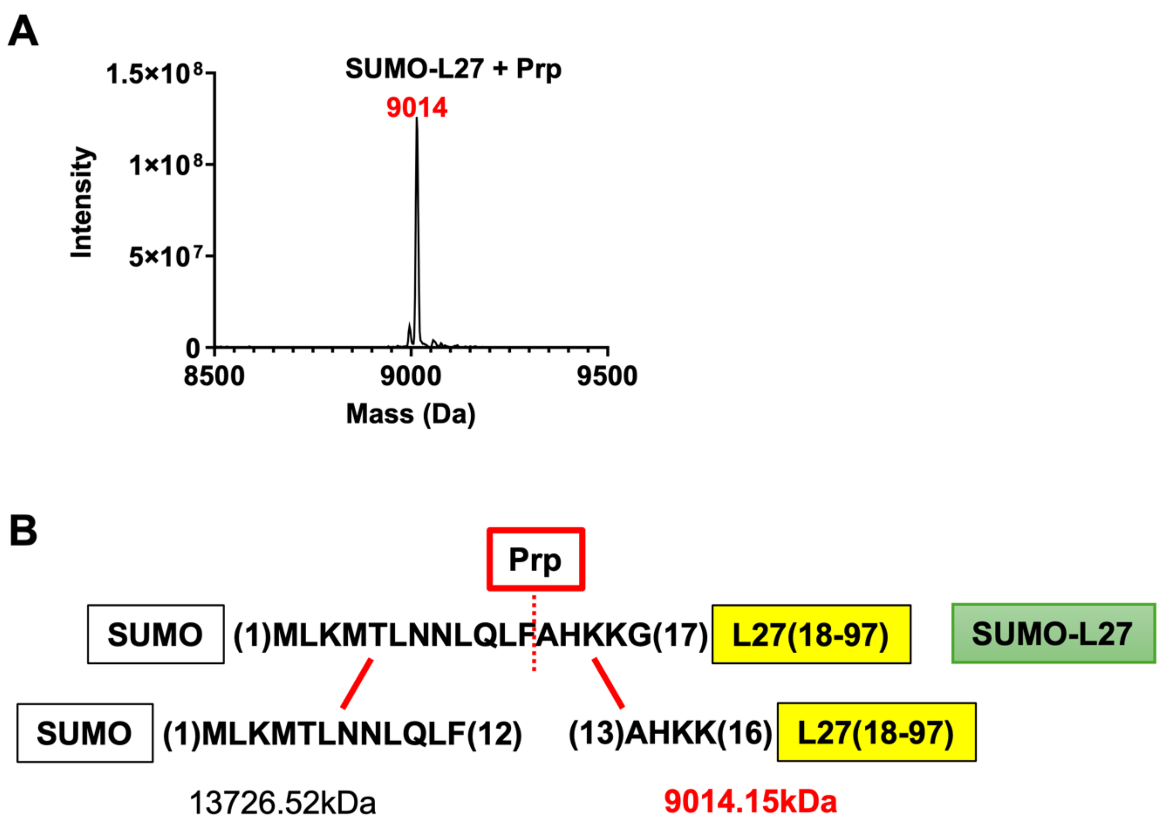


**Figure S6. Mass spectrometry of SUMO-L27 after Prp cleavage. (A)** Mass spectrum of the 8,500-9,500 Da range showing a single peak at 9,014 Da, corresponding to the cleaved L27ΔN. **(B)** Schematic diagram showing the fragments resulting from Prp cleavage, with theoretical masses listed.

### Figure S7:

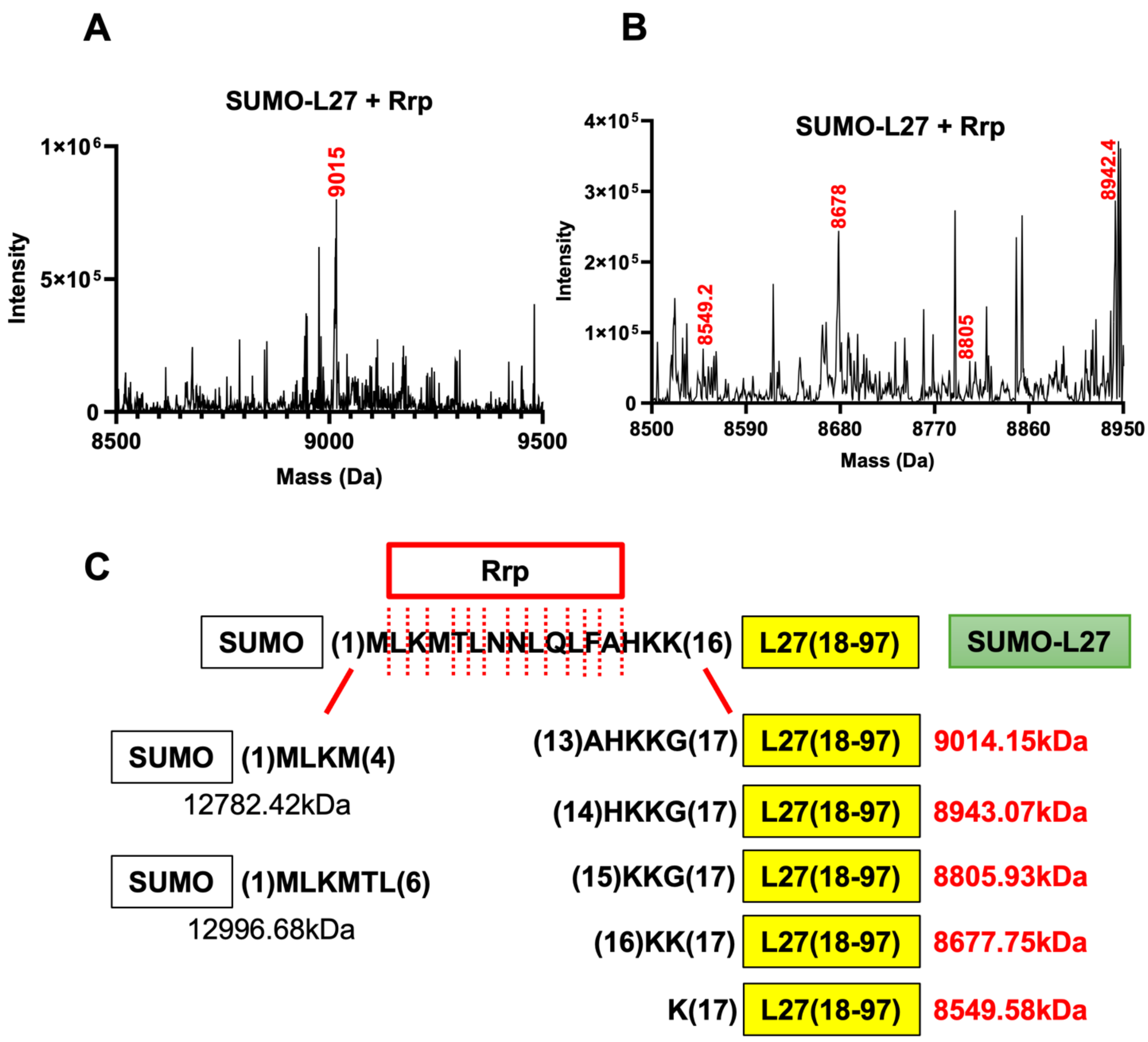


**Figure S7. Mass spectrometry of SUMO-L27 after Rrp cleavage. (A)** Mass spectrum of the 8,500-9,500 Da range showing a major peak at 9,015 Da and several minor peaks, indicating cleavage at multiple sites. **(B)** Mass spectrum showing only the range from 8,500-8,950 Da, indicating several peaks corresponding to cleaved SUMO-L27. **(C)** Schematic diagram showing the fragments resulting from Rrp cleavage, with theoretical masses listed.

#### Figure S8:

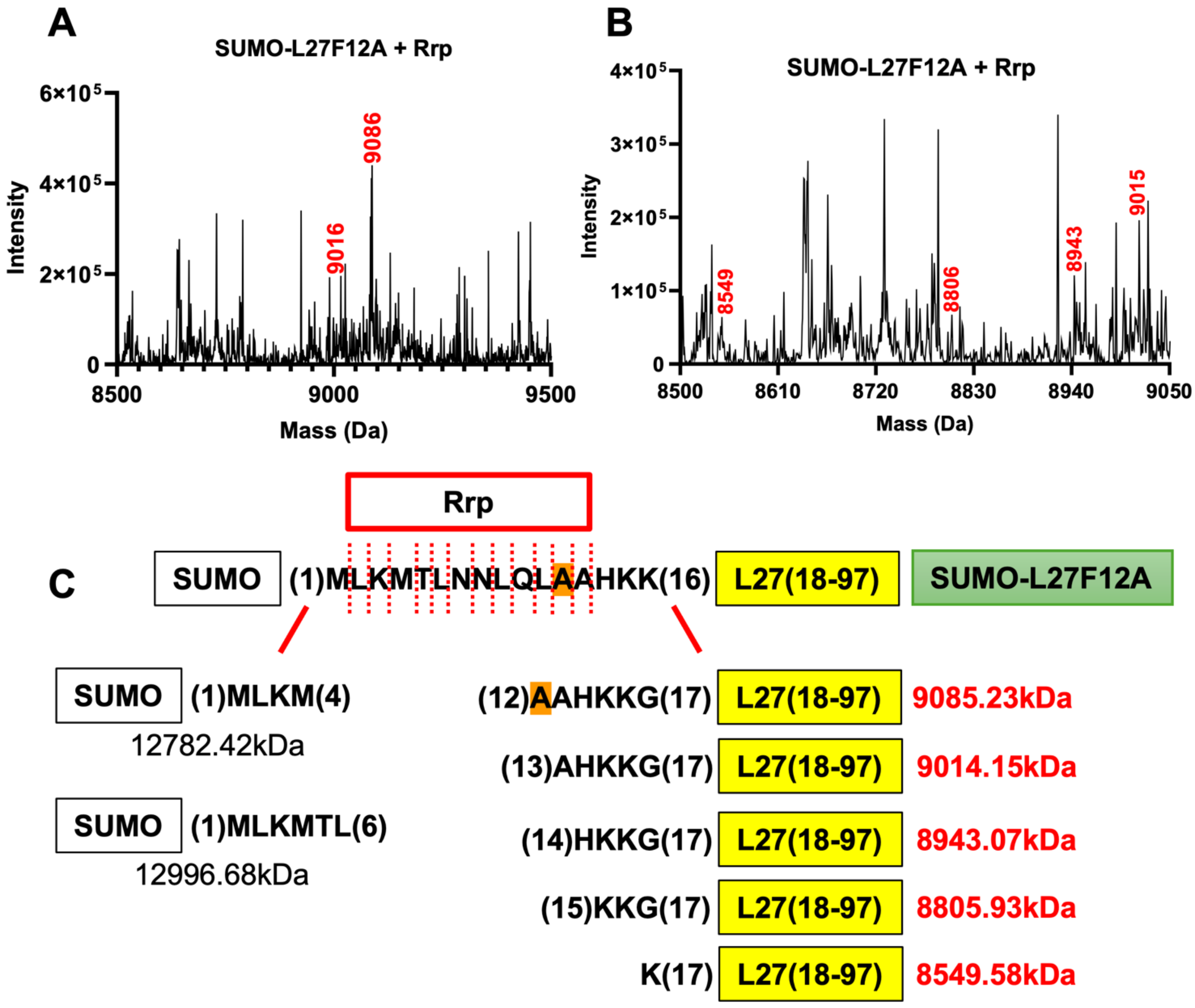


**Figure S8. Mass spectrometry of SUMO-L27F12A after Rrp cleavage. (A)** Mass spectrum of the 8,500-9,500 Da range. Several peaks are present, indicating cleavage at multiple sites. Two peaks corresponding to SUMO-L27F12A fragments at 9,016 and 9,086 Da are indicated. **(B)** Mass spectrum showing only the range from 8,500-8,950 Da. Some peaks corresponding to cleaved SUMO-L27F12A are indicated, but there are many additional peaks. **(C)** Schematic diagram showing the fragments resulting from cleavage of SUMO-L27F12A by Rrp, with theoretical masses listed.

**Table S1. Mass spectrometry of bL27.** Number of counts of all bL27 peptides identified by mass spectrometry in ribosomes isolated from TIGR4 wildtype and TIGR4 Δ*prp* strains after trypsin digestion of SDS-PAGE separated proteins. The N-terminal extension of bL27 (MLKMTLNNLQLF) was not detected.

| **L27 Peptides** | **Total Spectrum Count** | |
|---|---|---|
| | **TIGR4 wildtype** | **TIGR4 Δprp** |
| AADGQTVTGGSILYR | 22 | 28 |
| DDTLFAK | 3 | 5 |
| DTLFAK | 3 | 2 |
| GDDTLFAK | 5 | 3 |
| GGDDTLFAK | 35 | 27 |
| GGGSTSNGRDSQAK | 1 | 0 |
| GQTVTGGSILYR | 0 | 1 |
| GTHIYPGVNVGR | 26 | 27 |
| GVNVGR | 3 | 3 |
| KGGGSTSNGRDSQAK | 1 | 0 |
| MTLNNLQLFAHK | 0 | 1 |
| PGVNVGR | 2 | 1 |
| QLFAHK | 1 | 0 |
| QRGTHIYPGVNVGR | 3 | 1 |
| QVSVYPIAK | 2 | 3 |
| VEGVVR | 31 | 64 |
| VNVGRGGDDTLFAK | 1 | 1 |

**Table S2. Proteins detected by MS.** Total spectrum count measured by mass spectrometry of purified ribosomes from TIGR4 wildtype and TIGR4 Δ*prp* strains after trypsin digestion of SDS-PAGE separated proteins. The data shows the presence of peptides corresponding to bL27 and SP1145 (Rrp) and the absence of Prp. L5 and L21 are included as references. (Protein threshold 95%, peptide threshold 80%, minimum number of peptides 2, total number of proteins found 497.)

| **Protein** | **Gene** | **Accession No** | **Total spectrum count** | |
|---|---|---|---|---|
| | | | TIGR4 WT | MN001 (TIGR4 Δ*prp*) |
| L5 | *rplE (SP_0221)* | AAK74401.1 | 837 | 758 |
| L21 | *rplU (SP_1105)* | AAK75216.1 | 89 | 45 |
| L27 | *rpmA (SP_1107)* | AAK75218.1 | 43 | 44 |
| Prp | *prp (SP_1106)* | AAK75217.1 | 0 | 0 |
| SP1145 (Rrp) | *SP_1145* | AAK75255.1 | 91 | 58 |

**Table S3. Predicted masses resulting from cleavage of SUMO-L27 and SUMO-L27F12A.** The cleavage site is indicated by the arrow. Masses detected experimentally are highlighted in red.

| **Cut site** | **MW (Dalton)** | |
|---|---|---|
| **L27 fragment (C-terminal)** | **SUMO-L27** | **SUMO-L27F12A** |
| MLKMTLNN↓LQLFAHKKGGGSTS … (23-97) | 9515.77 | 9439.68 |
| MLKMTLNNL↓QLFAHKKGGGSTS … | 9402.61 | 9326.52 |
| MLKMTLNNLQ↓LFAHKKGGGSTS .. | 9274.48 | 9198.39 |
| MLKMTLNNLQL↓FAHKKGGGSTS … | 9161.32 | **9085.23** |
| MLKMTLNNLQLF↓AHKKGGGSTS … | **9014.15** | **9014.15** |
| MLKMTLNNLQLFA↓HKKGGGSTS … | **8943.07** | **8943.07** |
| MLKMTLNNLQLFAH↓KKGGGSTS … | **8805.93** | **8805.93** |
| MLKMTLNNLQLFAHK↓KGGGSTS … | **8677.75** | 8677.75 |
| MLKMTLNNLQLFAHKK↓GGGSTS … | **8549.58** | **8549.58** |
| MLKMTLNNLQLFAHKKG↓GGSTS … | 8492.53 | 8492.53 |
| MLKMTLNNLQLFAHKKGG↓GSTS … | 8435.48 | 8435.48 |
| MLKMTLNNLQLFAHKKGGG↓STS … | 8378.42 | 8378.42 |
| MLKMTLNNLQLFAHKKGGGS↓TS … | 8291.35 | 8291.35 |
| MLKMTLNNLQLFAHKKGGGST↓S … | 8190.24 | 8190.24 |
| **SUMO-L27 fragment (N-terminal)** | **SUMO-L27** | **SUMO-L27F12A** |
| SUMO↓ | 12278.7 | 12278.7 |
| SUMO-M↓ | 12409.89 | 12409.89 |
| SUMO-ML↓ | 12523.05 | 12523.05 |
| SUMO-MLK↓ | 12651.23 | 12651.23 |
| SUMO-MLKM↓ | **12782.42** | **12782.42** |
| SUMO-MLKMT↓ | 12883.52 | 12883.52 |
| SUMO-MLKMTL↓ | **12996.68** | **12996.68** |
| SUMO-MLKMTLN↓ | 13110.79 | 13110.79 |
| SUMO-MLKMTLNN↓ | 13224.89 | 13224.89 |
| SUMO-MLKMTLNNL↓ | 13338.05 | 13338.05 |
| SUMO-MLKMTLNNLQ↓ | 13466.18 | 13466.18 |
| SUMO-MLKMTLNNLQL↓ | 13579.34 | 13579.34 |
| SUMO-MLKMTLNNLQLF↓ | **13726.52** | 13650.42 |
| SUMO-MLKMTLNNLQLFA↓ | 13797.6 | 13721.5 |
| SUMO-MLKMTLNNLQLFAH↓ | 13934.74 | 13858.64 |
| SUMO-MLKMTLNNLQLFAHK↓ | 14062.91 | 13986.81 |
| SUMO-MLKMTLNNLQLFAHKK↓ | 14191.09 | 14114.99 |
| SUMO-MLKMTLNNLQLFAHKKG↓ | 14248.14 | 14172.04 |
| SUMO-MLKMTLNNLQLFAHKKGG↓ | 14305.19 | 14229.09 |
| SUMO-MLKMTLNNLQLFAHKKGGG↓ | 14362.24 | 14286.14 |
| SUMO-MLKMTLNNLQLFAHKKGGGS↓ | 14449.32 | 14373.22 |
| SUMO-MLKMTLNNLQLFAHKKGGGST↓ | 14550.42 | 14474.33 |
| SUMO-MLKMTLNNLQLFAHKKGGGSTS↓ | 14637.5 | 14561.41 |

**Table S4. List of genes around *SP_1145* in *S. pneumoniae* TIGR4.**

| TIGR4 gene | D39 gene | Size (aa) | Annotated function | Homologs | PDB IDs | Prob* | Length* |
|---|---|---|---|---|---|---|---|
| *SP_1124* | *SPD_1008* | | glycogen synthase | | | | |
| *SP_1125* | *SPD_1009* | | hypothetical | | | | |
| *SP_1126* | *SPD_1011* | | hypothetical | | | | |
| *SP_1127* | *SPD_1010* | | hypothetical | | | | |
| *SP_1128* | *SPD_1012* | | enolase | | | | |
| *SP_1129* | – | 387 | phage integrase family | Integrase, transposase | 6EMY, 1Z1B | 100.0 | 303 |
| *SP_1130* | – | 282 | Transcriptional regulator | transcriptional regulator, SaPI Stl repressor | 7ZCV, 7P4A | 99.2 | 172 |
| *SP_1131* | – | 117 | Transcriptional regulator | ComR transcriptional regulator, quorum sensing | 7N1N, 5FD4 | 98.4 | 62 |
| *SP_1132* | – | 88 | hypothetical | Excisionase; repressor; terS | 8DGL, 6G1T, 7LW0 | 99.0 | 55 |
| *SP_1133* | – | 50 | hypothetical | none | none | <60 | – |
| *SP_1134* | – | 205 | hypothetical | DNA damage response protein DdrC | 7UDI | 99.4 | 173 |
| *SP_1135* | – | 150 | hypothetical | Ribosomal protein L29 | 8UXA | 93.1 | 76 |
| *SP_1136* | – | 148 | hypothetical | DNA replication protein DnaD, helicase loader | 8OJJ, 2ZC2, 2I5U | 97.2 | 102 |
| *SP_1137* | – | 294 | putative GTP binding | DnaI primase/helicase loader | 4M4W, 2QGZ | 99.9 | 187 |
| *SP_1138* | – | 36 | hypothetical | none | none | <60 | – |
| *SP_1139* | – | 216 | hypothetical | none | none | <60 | – |
| *SP_1140* | – | 142 | hypothetical | Small terminase TerS (phages PaP3, SF6) | 7JOQ, 6W7T | 99.3 | 96 |
| *SP_1141* | – | 103 | hypothetical | none | none | <60 | – |
| *SP_1142* | – | 136 | hypothetical | RNA polymerase sigma factor | 6IN7, 2H27 | 96.4 | 65 |
| *SP_1143* | – | 121 | hypothetical | HigB toxin, Toxin-Antitoxin complex, RNAse | 6AF4, 7AWK | 99.8 | 120 |
| *SP_1144* | – | 97 | hypothetical | HigA antitoxin, Toxin-antitoxin complex | 6AF4, 7AWK | 98.9 | 96 |
| *SP_1145* | – | 396 | hypothetical | Anbu ancestral proteasome beta, hydrolase | 5NYW, 5LE5 | 98.8 | 207 |
| *SP_1146* | – | 58 | hypothetical | none | none | <60 | – |
| *SP_1147* | *SPD_1013* | | Integrase/recombinase | | | | |
| *SP_1148* | – | | IS630-Spn1, Transposase orf2 | | | | |
| *SP_1149* | *SPD_1014* | | IS630-Spn1, Transposase orf1 | | | | |

*Prob = probability from HHpred; Length = number of amino acids of the matched region.
